# Cysteine-mediated bortezomib resistance is governed by α-ketoacid availability

**DOI:** 10.64898/2026.07.06.736827

**Authors:** Jennifer A. Brain, Sarah M. Chang, Maximilian Kobiesa, Leah G. Rector, Michele Ceribelli, Sky H. Kim, Zhaoqi Li, Brian T. Do, Kelli J. Che, David Holland, Samuel Block, Renee Chang, Gillian E. Oaks, Craig J. Thomas, Matthew G. Vander Heiden, Lucas B. Sullivan

## Abstract

Dissecting how metabolite availability impacts drug sensitivity is critical for understanding therapy resistance. While proteasome inhibitors like the boronic acid proteasome inhibitor bortezomib are cornerstones of therapy for multiple myeloma, the clinical utility of these drugs is often limited by development of resistance, the metabolic drivers of which remain poorly understood. In this study, we find that α-ketoacids, including pyruvate, increase sensitivity to boronic acid proteasome inhibitors independent of their conventional roles in metabolism. Instead, α-ketoacids directly react with intracellular cysteine to form thiazolidines, sequestering cysteine away from forming conjugates with boronic acid proteasome inhibitors and detoxifying these drugs. Preventing cysteine-mediated detoxification through α-ketoacid supplementation increases the efficacy of boronic acid proteasome inhibitors, leading to greater proteasome inhibition and cytotoxicity that is reversed by cysteine supplementation. These findings suggest that modulating available cysteine through α-ketoacid interactions can impact effective levels of some drugs in cells, and represent a potential strategy to overcome resistance and maximize the efficacy of boronic acid proteasome inhibitors.

## Introduction

Cancer cells exhibit altered cell metabolism to support rapid proliferation, survival under stress, and resistance to therapy. Of note, cell-extrinsic factors such as environmental metabolite levels can impact cancer cell metabolism, tumor growth, and drug sensitivity^1–5^, but how endogenous environmental metabolites might interact with therapeutic agents to shape responses has not been extensively studied. Several metabolites have emerged as key modulators of cancer cell drug sensitivity, including the amino acid cysteine (CYS). CYS plays a central role in cancer cell survival and redox control, serving as proteinogenic amino acid for protein synthesis and a key precursor for glutathione, a major intracellular antioxidant. CYS in cancer cells can be imported from the extracellular environment as cystine (CYS_2_) via the glutamate/CYS_2_ antiporter xCT and reduced to CYS intracellularly. Changes in xCT subunit expression or activity can directly impact intracellular CYS levels, with corresponding effects on resistance to chemotherapies that generate reactive oxygen species (ROS) or promote ferroptosis^6–8^.

Pyruvate, a key metabolite in central carbon metabolism, can also alter cell sensitivity to therapies through distinct metabolic roles. For instance, pyruvate can serve as an electron acceptor, regenerating NAD+ from NADH, causing resistance to mitochondrial respiration inhibitors^3,9,10^. Pyruvate-driven anaplerosis has also been associated with resistance to TCA cycle inhibition, kinase inhibition, and disruptions to glutamine metabolism^11–14^. Interestingly, pyruvate has recently been shown to alter sensitivity to the proteasome inhibitor bortezomib through supporting mitochondrial metabolism, regulating ROS production, and influencing proteostasis^15^. Collectively, these observations highlight that metabolites, including pyruvate, can impact drug therapy response.

Proteosome inhibitors represent a class of cancer drugs that are particularly useful for treatment of multiple myeloma (MM). The proteasome is a large protease complex responsible for the selective degradation of intracellular proteins, thereby maintaining protein homeostasis and regulating various cellular processes, including the cell cycle, gene expression, and antigen presentation^16,17^. Proteins targeted for degradation are first tagged with chains of ubiquitin, a small regulatory protein that can signal recognition by the proteasome. The proteasome consists of a 19S regulatory particle and a 20S catalytic core particle, the latter of which houses six proteolytically active β subunits: two copies each of the β1, β2, and β5 subunits^17,18^. The β5 subunit determines the rate of protein breakdown and is the subunit targeted by all clinically approved proteasome inhibitors^19,20^. Given the heightened dependency of some cancers on protein synthesis, particularly plasma cell malignancies including MM, targeting the proteasome has been effective for therapy. The first-in-class proteasome inhibitor bortezomib (Velcade^®^) was initially approved in 2003 for the treatment of relapsed and refractory MM and then subsequently approved for newly diagnosed MM and mantle cell lymphoma^21^. In 2012, carfilzomib (Kyprolis^®^) was approved for relapsed and refractory MM treatment^22^, and in 2015, the first oral PROTEASOME INHIBITOR ixazomib (Ninlaro^®^) was also approved for relapsed and refractory MM^23^. These clinically approved proteasome inhibitors fall into two distinct chemical classes: peptide boronates (bortezomib and ixazomib) and epoxyketones (carfilzomib).

Despite the clinical success of proteasome inhibitors, most patients develop resistance to these drugs leading to disease relapse. Some resistant cells acquire mutations in the PSMB5 β5 subunit to reduce drug binding, though these are relatively rare^24,25^. Additionally, resistance can be conferred by increased drug efflux or compensation by upregulating alternative protein degradation pathways^26–28^. Metabolic rewiring and perturbations can also contribute to PROTEASOME INHIBITOR resistance^29–33^. In many cases drug resistance is not understood, and identifying factors that modulate PROTEASOME INHIBITOR efficacy could help improve the use of these drugs.

In this study, we uncovered a novel interactions between metabolite availability and cancer cell drug sensitivity. Specifically, we find that pyruvate, levels of which can vary across tumor microenvironments^34^. increases cancer cell sensitivity to boronic acid proteasome inhibitors but not to other classes of proteasome inhibitors. Furthermore, we find that other α-ketoacids can similarly increase cancer cell sensitivity to boronic acid proteasome inhibitors. This increased cell sensitivity was not dependent on changes to cell redox state or central carbon metabolism. Instead, we found that α-ketoacids react with the amino acid cysteine, sequestering cysteine that could otherwise react with boronic acid groups of proteasome inhibitors and diminish their activity. Overall, we uncover interactions between boronic acid moieties and the endogenous metabolites cysteine, as well as between cysteine and α-ketoacid such as pyruvate, that to alter effective drug concentrations in cells. Taken together, these findings reveal a novel modulator of cancer cell sensitivity to boronic acid proteasome inhibitors by mitigating cysteine-dependent detoxification and, more broadly, demonstrate how chemical interactions between metabolites can modulate drug sensitivity and resistance.

## Results

### Pyruvate increases sensitivity to boronic acid proteasome inhibitors by enhancing proteasome inhibition

To investigate how pyruvate availability alters therapeutic sensitivity, we conducted a small molecule screen assessing A549 non-small cell lung cancer cells cultured in media with and without pyruvate (5 mM). We treated cells with the NCATS Mechanism Interrogation PlatEs (MIPE) library version 5.0, which includes 2480 oncology focused, mechanistically annotated agents that cover more than 800 distinct mechanisms of action, and measured viability in cells using CellTiter-Glo after 48 hours of exposure to a range of drug concentrations (**Figure 1A**)^35^. This small molecule screen found that cells were more resistant to multiple glutaminase inhibitors in the presence of pyruvate (**Extended Data Figure 1**), consistent with pyruvate supporting TCA cycle anaplerosis in glutamate-limiting conditions^11,14^. Additionally, we found that pyruvate increases sensitivity to the proteasome inhibitor bortezomib, consistent with recent observations^15^. Interestingly however, we observe that while some proteasome inhibitors, including bortezomib, were among the drugs most sensitized by the addition of pyruvate, other proteasome inhibitors were unaffected by pyruvate supplementation (**Figure 1B**). Considering the shared molecular target of proteasome inhibitors, we rationalized that the differences in cell viability in the presence of pyruvate might be explained by different chemical features of the small molecule proteosome inhibitors. Indeed, we observe that only proteasome inhibitors with a boronic acid functional group were sensitized by pyruvate supplementation (**Figure 1B**). To confrim this finding, we exposed A549 cells to the FDA-approved proteasome inhibitors bortezomib, ixazomib, and carfilzomib, with or without 1 mM pyruvate supplementation. Consistently, pyruvate increased cell sensitivity to boronic acid proteasome inhibitors, including bortezomib and ixazomib, while having no impact on cell sensitivity to carfilzomib (**Figure 1C-D, Extended Data Figure 2A**). We then evaluated the dose responsiveness of this effect by treating A549 cells with bortezomib and increasing doses of pyruvate, finding that pyruvate dose-dependently sensitizes cells to bortezomib (**Extended Data Figure 2B**). To further evaluate the ability of pyruvate to sensitize cells to boronic acid containing proteasome inhibitors, we treated A549 cells with the peptide aldehyde proteasome inhibitor MG-132 and its boronic acid containing variant MG-262. Pyruvate supplementation sensitized cells to MG-262, but not to MG-132, suggesting that the ability of pyruvate to increase sensitivity to proteasome inhibitors is specific to the presence of boronic acid (**Extended Data Figure 2C**). Because bortezomib and carfilzomib are approved for the treatment of multiple myeloma, we treated L-363 and AMO-1 multiple myeloma cells with either bortezomib or carfilzomib, with or without pyruvate. Consistent with results in A549 cells, pyruvate also sensitizes multiple myeloma cells to bortezomib but not carfilzomib (**Extended Data Figure 2D,E**). We observed similar effects of pyruvate on proteosome inhibitor sensitivity in the bile duct cancer cells CCLP1, the pancreatic adenocarcinoma cells SUIT2, and the chronic myeloid leukemia cells K562, indicating that pyruvate sensitization of cancer cells to boronic acid proteasome inhibitors is neither cell line- nor cancer type-specific (**Extended Data Figure 2F-H**).

**Figure 1.**
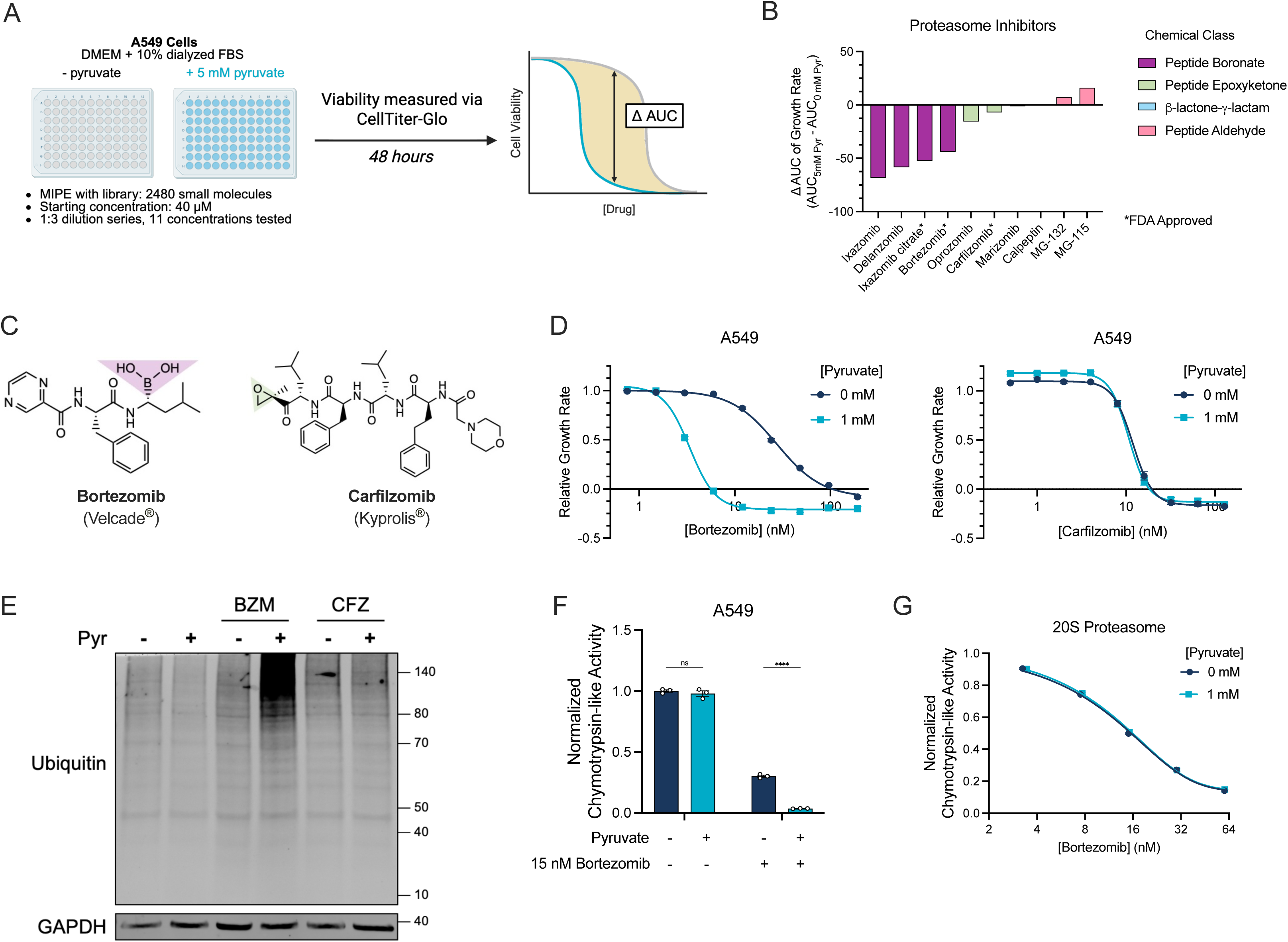
Pyruvate increases cell sensitivity to bortezomib and enhances proteasome inhibition of bortezomib-treated cells. **(A)** Schematic depicting the high-throughput quantitative small molecule screen using the MIPEv5 library. A549 cells were treated with the MIPEv5 library with and without 5 mM pyruvate for 48 hours. Cell viability was measured with Cell-Titer Glo, and relative, growth-rate corrected luminescence units were used to calculate area under the curve (AUC) values for each drug. For analysis of sensitivity to pyruvate, the difference in the area under the curve (*Δ*AUC) between conditions with or without pyruvate was calculated. **(B)** *Δ*AUC values between conditions with or without pyruvate for all proteasome inhibitors included in the MIPEv5 library and the chemical class of proteasome inhibitors. * indicates proteasome inhibitors that are currently FDA-approved. Drugs are colored by chemical class of their functional group **(C)** Chemical structures of bortezomib and carfilzomib. The functional group is highlighted by chemical class in (B). **(D)** Relative growth rate of A549 cells treated with or without 1 mM pyruvate and cotreated with increasing doses of bortezomib or carfilzomib for 96 hours. Growth rates are relative to the untreated conditions for each treatment condition, n=6 per condition. **(E)** Immunoblot for total ubiquitinated protein of A549 cells treated with and without 1 mM pyruvate, 15 nM bortezomib, or 15 nM carfilzomib for 24 hours as indicated. GAPDH is shown as a loading control. Immunoblot is a representative image of three independent biological replicates. **(F)** Cell-based proteasome activity assay for chymotrypsin-like activity in A549 cells treated with or without 1 mM pyruvate or 15 nM bortezomib for 6 hours, as indicated. Values were normalized to untreated conditions for each treatment condition and represent means, n=3 per condition. **(G)** Chymotrypsin-like activity of purified 20S proteasomes incubated with increasing doses of bortezomib with or without 1 mM pyruvate for 1 hour. Values represent means, n=3 per condition. For all panels values represent means, error bars are SEM. Relative growth rate curves were fitted with a nonlinear variable slope (four parameters) regression model. Statistical significance was assessed by two-way ANOVA with Sidak’s multiple comparison test (F). ns = not significant, ****p<0.0001.

We next asked whether pyruvate increased cell sensitivity to boronic acid proteasome inhibitors by enhancing the degree of proteasome inhibition. We first measured the accumulation of ubiquitinated proteins in A549 cells treated with low doses of either bortezomib or carfilzomib, with or without 1 mM pyruvate, for 24 hours. We observed that pyruvate increases the accumulation of ubiquitinated proteins in cells treated with bortezomib, while there was no change in ubiquitinated protein accumulation in cells treated with carfilzomib with or without pyruvate (**Figure 1E**). We next directly measured proteasome activity from cells treated with bortezomib and/or pyruvate by luminogenic substrates that specifically quantifies the activity of the rate-limiting chymotrypsin-like (β5) subunit of the 20S proteasome that is targeted by the majority of proteasome inhibitors. We find that pyruvate enhances inhibition of the chymotrypsin-like subunit at a given dose of bortezomib, but has no differential impact on proteasome inhibition by carfilzomib (**Figure 1F, Extended Data Figure 3A**). These data suggest that pyruvate elevates cell sensitivity to bortezomib by enhancing proteasome inhibition. We also employed a UbG76V-GFP reporter system to quantify ubiquitin-proteasome dependent proteolysis in AMO-1 multiple myeloma cells (AMO-1 UbGFP). We treated cells with either bortezomib or carfilzomib, with or without pyruvate^36^. We observed an increase in GFP signal in viable cells treated with both bortezomib and pyruvate compared to cells treated with bortezomib alone, but did not observe a change in UbGFP signal in cells treated with carfilzomib and pyruvate (**Extended Data Figure 3B-C**). Consistently, cells treated with both bortezomib and pyruvate exhibited greater cytotoxicity that cells treated with bortezomib alone (**Extended Data Figure 3D-E**). Lastly, we tested the effects of pyruvate on proteasome inhibition from bortezomib using purified 20S proteasomes to evaluate whether pyruvate directly sensitizes the proteosome to bortezomib^37^. Interestingly, there was no change in the chymotrypsin-like activity of purified 20S proteasomes treated with bortezomib in the presence of pyruvate, indicating that pyruvate does not directly modify the proteasome or bortezomib to explain how it impacts the extent of proteosome inhibition by boron-containing proteosome inhibitors in cells (**Figure 1G**).

### Pyruvate does not increase cell sensitivity to bortezomib through its canonical metabolic processes

Proteasome activity can be impacted by both mitochondrial metabolism and the NAD+/NADH ratio^15,30,38^. Since pyruvate treatment may mediate its effects on bortezomib sensitivity through these effects on metabolism, including serving as an electron acceptor to increase the cell NAD+/NADH ratio, we sought to evaluate treatments that could uncouple pyruvate abundance from its effects on NAD+/NADH (**Figure 2A**)^10^. First, we evaluated how lactate treatment influenced bortezomib sensitivity, as high doses of lactate impair pyruvate reduction by Lactate Dehydrogenase (LDH), increasing pyruvate abundance while lowering the NAD+/NADH ratio^39^. Indeed, lactate treatment was sufficient to decrease the NAD+/NADH ratio and increase pyruvate levels (**Figure 2B-C**). Notably, lactate also sensitized cells to bortezomib to a degree that was commensurate with the increase in pyruvate, suggesting the lactate sensitizing effect is mediated by increasing pyruvate rather than by modulating the cell NAD+/NADH ratio (**Figure 2B-D**). Moreover, lactate supplementation did not further increase cell sensitivity to bortezomib in cell culture media containing pyruvate, also suggesting that any effects of lactate on increasing cell sensitivity to bortezomib are caused by changing pyruvate levels (**Extended Data Figure 4A**).

**Figure 2.**
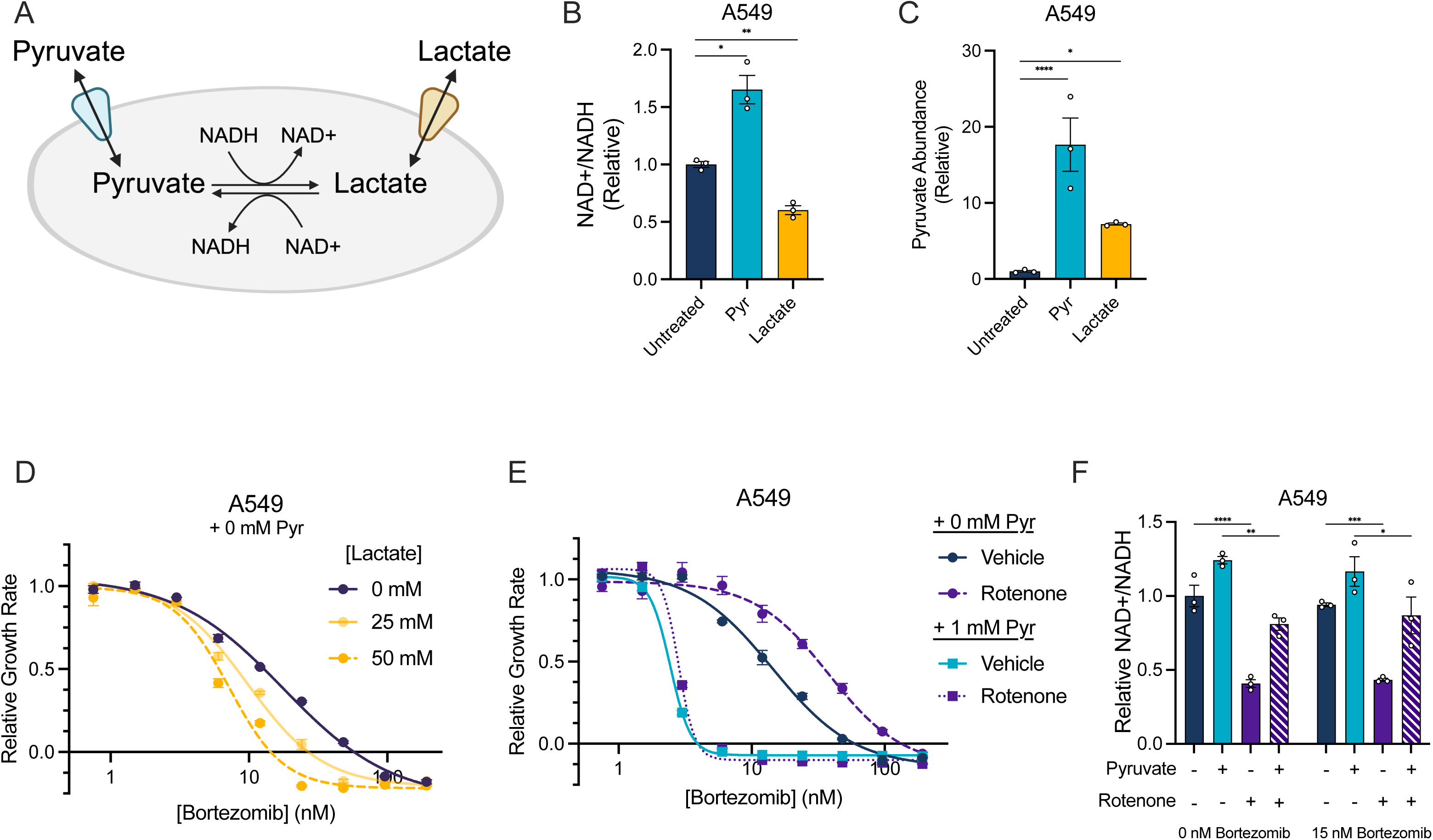
Pyruvate does not increase cell sensitivity to bortezomib by changing the cell NAD+/NADH ratio. **(A)** Schematic depicting the impact of pyruvate or lactate cellular import on the cell NAD+/NADH ratio. Pyruvate supplementation increases the NAD+/NADH ratio whereas lactate supplementation decreases the NAD+/NADH ratio. **(B)** The relative NAD+/NADH ratio as measured by LC-MS from A549 cells treated with 1 mM pyruvate or 50 mM lactate for 1.5 hours. Ratios are relative to untreated cells, n=3 per condition. **(C)** Relative abundance of intracellular pyruvate as measured by LC-MS from A549 cells upon treatment with 1 mM pyruvate or 50 mM lactate for 1.5 hours. Abundances are relative ion counts to untreated cells, n=3 per condition. **(D)** Relative growth rate of A549 cells treated with or without 25 mM or 50 mM lactate and cotreated with increasing doses of bortezomib for 96 hours. Growth rates are relative to the untreated condition, n=3 per condition. **(E)** Relative growth rate of A549 cells treated with or without 50 nM rotenone or 1 mM pyruvate cotreated with increasing doses of bortezomib for 96 hours. Growth rates are relative to the untreated condition, n=6 per condition. **(F)** The NAD+/NADH ratio measured via enzymatic assay of A549 cells treated with 1 mM pyruvate, 50 nM rotenone, and 15 nM bortezomib as indicated for 24 hours. Values represent means, n=3 per condition. For all panels values represent means, error bars are SEM. Relative growth rate curves were fitted with a nonlinear variable slope (four parameters) regression model. Statistical significance was assessed by an unpaired two-tailed student’s t-test with a bonferroni correction (B-C) or with a two-way ANOVA with Sidak’s correction for multiple comparisons (F). *p < 0.05, **p < 0.01, ***p < 0.001, ****p < 0.0001.

We next sought to determine if pyruvate supplementation increased cell sensitivity to bortezomib by affecting mitochondrial metabolism. First, we co-treated cells with pyruvate and the mitochondrial electron transport chain complex I inhibitor rotenone, which has been found to confer resistance to proteasome inhibitors^30^. As expected, cells treated with rotenone were moderately growth-impaired, and this effect was partially rescued by pyruvate (**Extended Data Figure 4B**)^9,10^. While rotenone decreased the cell NAD+/NADH ratio in cells treated with pyruvate, and reduced sensitivity to bortezomib in the absence of pyruvate, rotenone did not alter cell sensitivity to bortezomib in the presence of pyruvate (**Figure 2E-F**). Additionally, co-treating cells with the pyruvate dehydrogenase kinase inhibitor dichloroacetic acid (DCA), which stimulates pyruvate-driven mitochondrial metabolism via disinhibiting pyruvate dehydrogenase, did not impact cell sensitivity to bortezomib in media with or without pyruvate (**Extended Data Figure 4C**). Furthermore, decreasing mitochondrial pyruvate uptake with the mitochondrial pyruvate carrier inhibitor UK5099 also did not alter cell sensitivity to bortezomib (**Extended Data Figure 4D**). Together, these data highlight that pyruvate increases cell sensitivity to bortezomib by enhancing proteasome inhibition independent of an impact on mitochondrial metabolism or the cell NAD+/NADH ratio.

### Diverse α-ketoacid molecules redundantly promote bortezomib sensitivity

Given that several of the canonical metabolic effects of pyruvate did not alter cell sensitivity to bortezomib, we next asked if its chemical features might be responsible for promoting cell sensitivity to boronic acid containing proteasome inhibitors. Pyruvate is an α-ketoacid, so we considered whether other α-ketoacids could similarly impact bortezomib sensitivity. We first treated A549 cells with α-ketobutyrate (αKB) and observed an increase in sensitivity to bortezomib (**Extended Data Figure 5A-B**). However, αKB treatment also increased intracellular pyruvate, making it impossible to discern if the effects of αKB were direct or mediated by the increase in pyruvate (**Extended Data Figure 5C**). Next, we treated cells with α-ketoacids that are not expected to alter intracellular pyruvate levels, including the TCA cycle metabolite α-ketoglutarate (αKG) and the phenylalanine metabolism intermediate phenylpyruvate. Both αKG and phenylpyruvate robustly increased sensitivity to bortezomib or ixazomib while only having minimal effects on intracellular pyruvate (**Figure 3A-D, Extended Data Figure 5D**). Notably, the sensitization effect of αKG and phenylpyruvate was much greater than that observed from lactate (**Figure 2D**) while having a smaller effect on pyruvate levels, indicating that these α-ketoacids likely phenocopy the bortezomib sensitivity independent of any effect on pyruvate levels. Moreover, neither αKB, αKG, nor phenylpyruvate further sensitized cells to boronic acid proteasome inhibitors in media containing pyruvate (**Extended Data Figure 5B, E-F**). Taken together, these data suggest that the mechanism by which pyruvate sensitizes cells to boronic acid proteasome inhibitors is phenocopied by other α-ketoacids, indicating that their shared effects may be dependent on their overlapping chemical properties rather than their metabolic functions.

**Figure 3.**
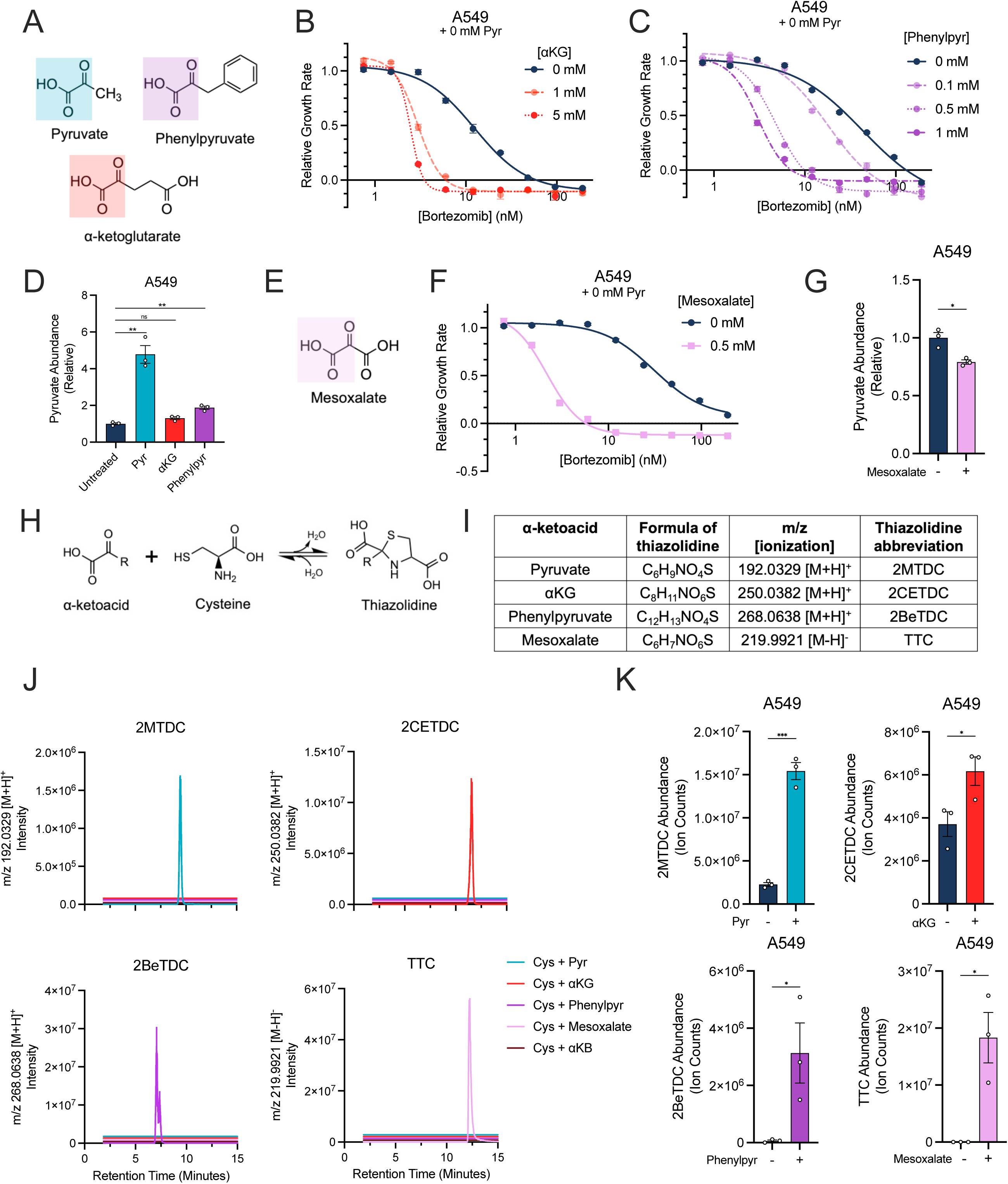
α-ketoacids increase cell sensitivity to bortezomib and form thiazolidines with cysteine. **(A)** Chemical structures of the α-ketoacids pyruvate (pyr), α-ketoglutarate (αKG), and phenylpyruvate (phenylpyr) with the α-ketoacid highlighted in blue, red, or purple, respectively. **(B)** Relative growth rate of A549 cells treated with or without 1 mM or 5 mM αKG and cotreated with increasing doses of bortezomib for 96 hours. Growth rates are relative to the untreated condition, n=6 per condition. **(C)** Relative growth rate of A549 cells treated with or without 0.1 mM, 0.5 mM or 1 mM phenylpyruvate and cotreated with increasing doses of bortezomib for 96 hours. Growth rates are relative to untreated condition, n=6 per condition. **(D)** Relative abundance of intracellular pyruvate as measured by LC-MS from A549 cells treated with 1 mM pyruvate, 5 mM αKG, or 1 mM phenylpyruvate for 1.5 hours. Abundances are relative ion counts to untreated cells, n=3 per condition. **(E)** Chemical structure of the α-ketoacid Mesoxalate with the α-ketoacid highlighted in pink. **(F)** Relative growth rate of A549 cells treated with or without 0.5 mM mesoxalate and cotreated with increasing doses of bortezomib. Growth rates are relative to untreated condition, n=3 per condition. **(G)** Relative abundance of intracellular pyruvate as measured by LC-MS from A549 cells treated with 0.5 mM mesoxalate for 1.5 hours. Abundances are relative ion counts to untreated cells, n=3 per condition. **(H)** Schematic depicting mechanism by which α-ketoacids can react with cysteine (CYS). **(I)** Table of predicted thiazolidine between cysteine and the α-ketoacids pyruvate, αKG, phenylpyruvate, or Mesoxalate, their chemical formulas, their expected m/z and ionization, and their abbreviations: CYS-αKG <u>2</u>-(2-<u>c</u>arboxy<u>e</u>thyl)-1,3-thiazolidine-2,4-<u>dic</u>arboxylate (2CETDC), CYS-phenylpyruvate <u>2</u>-<u>be</u>nzyl-1,3-thiazolidine-2,4-<u>dic</u>arboxylate (2BeTDC), (CYS-mesoxalate 1,3-thiazolidine-2,2,4- tri<u>c</u>arboxylate (TTC). **(J)** LC-MS ion chromatograms filtered by m/z for each predicted thiazolidine filtered by m/z. Purified chemical standards of CYS were incubated with or without each of the α-ketoacids overnight at 4**°**C. **(K)** LC-MS ion counts of cells treated with or without 1 mM Pyr, 5 mM αKG, 1 mM phenylpyruvate, or 0.5 mM Mesoxalate for 1.5 hours. n=3 per condition. For all panels values represent means, error bars are SEM. Relative growth rate curves were fitted with a nonlinear variable slope (four parameters) regression model. Statistical significance was assessed with an unpaired two-tailed student’s t-test with a bonferroni correction (D), or with an unpaired two-tailed student’s t-test (G, K). ns=not significant, *p < 0.05, **p < 0.01, ***p < 0.001, ****p < 0.0001.

To further separate the sensitization effects of α-ketoacids on boronic acid proteasome inhibitor sensitivity from any potential shared metabolic effects, we tested the effects of mesoxalate, a synthetic α-ketoacid molecule not known to occur in or function in human metabolism (**Figure 3E)**, on bortezomib sensitivity. Notably, mesoxalate increased cell sensitivity to bortezomib and ixazomib without increasing intracellular pyruvate levels (**Figure 3F-G, Extended Data Figure 5G**). Mesoxalate treatment also did not impact cell sensitivity to bortezomib or ixazomib in the presence of pyruvate, suggesting a redundant mechanism of sensitization (**Extended Data Figure 5G-H**). Importantly, neither phenylpyruvate nor mesoxalate sensitized cells to carfilzomib (**Extended Data Figure 5I-J**). Together, these data suggest that α-ketoacids, irrespective of any metabolic roles, increase cell sensitivity to boronic acid proteasome inhibitors.

### Bortezomib sensitizing α-ketoacids form adducts with CYS

Prior studies have noted that supplementation with the amino acid cysteine (CYS) or its derivative glutathione could suppress bortezomib-induced cytotoxicity^33,40^. We and others have found that the carbonyl of pyruvate can react with intracellular CYS to reversibly form the thiazolidine compound 2-methyl-2,4-thiazolidine dicarboxylate (2MTDC), which functionally decreases free cysteine availability (**Figure 3H-I**)^41–44^. We thus considered the possibility that these α-ketoacid molecules may impact boronic acid proteasome inhibitor sensitivity by sequestering CYS. First, we evaluated if bortezomib sensitizing α-ketoacid metabolites could form their respective predicted thiazolidine products by combining purified CYS and α-ketoacids in solution and measuring the expected thiazolidine m/z products by liquid chromatography-mass spectrometry (LC-MS) (**Figure 3I**). Indeed, this approach yielded product peaks with m/z values that corresponded to the predicted thiazolidine fate for pyruvate, αKG, phenylpyruvate, mesoxalate, and αKB (**Figure 3I-J, Extended Data Figure 6A-B**). Importantly, each of the thiazolidine products was only detected in samples containing both CYS and its corresponding parent α-ketoacid. We will refer to each of the CYS-α-ketoacid thiazolidines as follows: CYS-αKG <u>2</u>-(2-<u>c</u>arboxy<u>e</u>thyl)-1,3-thiazolidine-2,4-<u>dic</u>arboxylate (2CETDC), CYS-phenylpyruvate <u>2</u>-<u>be</u>nzyl- 1,3-thiazolidine-2,4-<u>dic</u>arboxylate (2BeTDC), (CYS-mesoxalate 1,3-thiazolidine-2,2,4- tri<u>c</u>arboxylate (TTC), CYS-αKB <u>2</u>-<u>e</u>thyl-1,3-thiazolidine-2,4-<u>dic</u>arboxylate (2ETDC).

To evaluate if thiazolidine formation could explain the impact of α-ketoacid supplementation on cell sensitivity to boronic acid proteasome inhibitors, we next determined if they form in cells upon α-ketoacid metabolite treatments. Indeed, upon treating cells with each α-ketoacid, we detected increased thiazolidine product abundances corresponding to each α-ketoacid treatment condition (**Figure 3K, Extended Data Figure 6C**). As expected, each α-ketoacid also increased 2MTDC abundance proportional to each α-ketoacid effect on intracellular pyruvate (**Figure 3D, G, Extended Data Figure 5C, Extended Data Figure 6D)**. We next confirmed that these newly described thiazolidine molecules formed through reversible reactions with CYS as occurs with other CYS-derived thiazolidine metabolites^42^. Thus, we extracted cells with or without the irreversible thiol conjugating agent N-ethylmaleimide (NEM), which conjugates free cysteine and shifts the equilibrium of thiazolidines towards disassociation and thereby depleting thiazolidine levels. Indeed, intracellular levels of the thiazolidine molecule formed between CYS and each α-ketoacid were decreased upon extraction in NEM compared to standard extractions without NEM (**Extended Data Figure 6E**). To confirm the shared identity between the synthesized thiazolidines and those in α-ketoacid treated A549 cell extracts we performed MS/MS fragmentation for each, finding strong overlap in fragmentation patterns for 2MTDC, 2CETDC, 2BeTDC, 2ETDC, and TTC (**Extended Data Figure 7A-E)**.

Because α-ketoacids can react with CYS intracellularly, we next evaluated if α-ketoacid treatment decreases intracellular CYS levels. LC-MS based measurements of CYS requires thiol conjugation (e.g. by NEM) to mitigate spontaneous oxidation and mixed disulfide generation that otherwise obscures quantification. However, because of thiazolidine reversibility during NEM extraction, this approach also overestimates the abundance of free CYS in cells. Nonetheless, we rationalized that because the depletion of thiazolidine metabolites upon NEM extraction was not complete (**Extended Data Figure 6E**), we could still qualitatively assses if CYS pools were impacted by α-ketoacid treatment by measuring CYS-NEM. Indeed, with most α-ketoacid treatments, we observed a decrease in CYS-NEM, corroborating that α-ketoacids can decrease free intracellular CYS abundance (**Extended Data Figure 7F**). Together, these results indicate that α-ketoacids react with free CYS to sequester the CYS as thiazolidine metabolites. They also suggest that the resulting decrease in free CYS might explain why α-ketoacids increase cell sensitivity to boronic acid containing proteasome inhibitors.

### Cysteine reacts with boronic acid proteasome inhibitors to diminish their efficacy

If α-ketoacid treatment increases cell sensitivity to bortezomib via decreased CYS availability, we hypothesized that increasing intracellular CYS would promote bortezomib resistance. To alter intracellular CYS abundance we cultured cells in standard conditions (200 μM CYS_2_) and high CYS_2_-containing media (700 μM CYS_2_), which caused corresponding effects on intracellular CYS abundance (**Figure 4A**), and then evaluated sensitivity to proteasome inhibitors in each media context. Notably, culture in high CYS_2_ media without pyruvate promoted cell resistance to bortezomib and ixazomib (**Figure 4B, Extended Data Figure 8A**). This resistance in high CYS_2_ media was suppressed upon supplementation with pyruvate (**Extended Data Figure 8B-C**). Importantly, sensitivity to carfilzomib was unaffected by high CYS_2_ conditions (**Figure 4C**). These data indicate that, as with α-ketoacids treatments, the sensitivity shifting effects of CYS are dependent on the presence of boronic acid in drug function.

**Figure 4.**
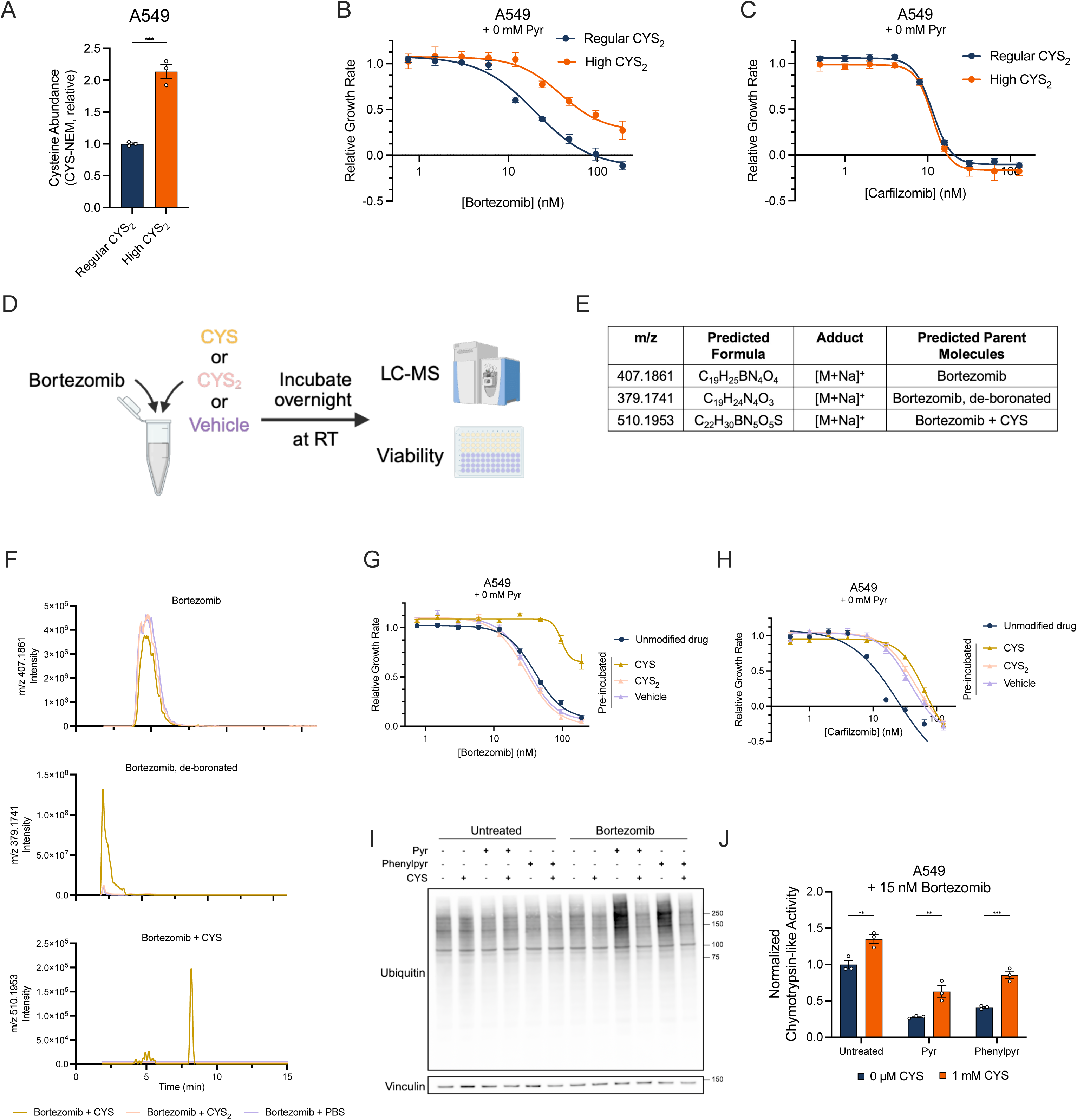
Exogenous cysteine supplementation reverses the increased cell sensitivity and proteasome inhibition by α-ketoacids in bortezomib-treated cells. **(A)** Relative abundance of total intracellular CYS, measured as CYS-NEM, from A549 cells treated with 200 μM CYS_2_ or 700 μM CYS_2_ for 1.5 hours. Abundances are relative ion counts to regular (200 μM) CYS_2_ cells. n=3 replicate wells per condition. **(B)** Relative growth rate of A549 cells upon treatment with 200 μM CYS_2_ or 700 μM CYS_2_ and cotreatment with varying doses of bortezomib as measured by SRB. Growth rates are relative to the 0 nM bortezomib treated wells for each treatment condition. n=6 replicate wells per condition. **(C)** Relative growth rate of A549 cells upon treatment with 200 μM CYS_2_ or 700 μM CYS_2_ and cotreatment with varying doses of carfilzomib as measured by SRB. Growth rates are relative to the 0 nM carfilzomib treated wells for each treatment condition. n=6 replicate wells per condition. **(D)** Schematic depicting bortezomib reacting with PBS or CYS in cell-free reactions. **(E)** Table of bortezomib-based LC-MS sodium ions including their m/z and the predicted formula and adduct. **(F)** LC-MS chromatograms show detection of all three bortezomib sodium adducts in samples incubated in PBS or CYS or CYS_2_ overnight at 4**°**C. **(G)** Relative growth rate of A549 cells upon treatment with unmodified bortezomib or bortezomib that had been previously incubated with CYS, CYS_2_, or PBS overnight at 4**°**C as measured by SRB. Growth rates are relative to the 0 nM bortezomib-treated wells for each treatment condition. n=6 replicate wells per condition. **(H)** Relative growth rate of A549 cells upon treatment with unmodified carfilzomib or carfilzomib that had been previously incubated with CYS overnight at 4**°**C as measured by SRB. Growth rates are relative to the 0 nM carfilzomib-treated wells for each treatment condition. n=6 replicate wells per condition. **(I)** Immunoblot for total ubiquitinated protein of A549 cells treated with and without 1 mM pyruvate, 1mM phenylpyruvate, 15 nM bortezomib, and 1 mM cysteine, as indicated. Vinculin is shown as a loading control. **(J)** Chymotrypsin-like proteasome activity was measured in A549 cells treated with 1 mM pyruvate or 1 mM phenylpyruvate with and without supplementation of 1 mM cysteine in the presence of 15 nM bortezomib. Activity is relative to untreated 0 μM CYS condition. n=3 technical replicates. For all panels, values represent means, error bars are SEM. Relative growth rate curves were fitted with a nonlinear variable slope (four parameters) regression model. Statistical significance was assessed with a student’s t-test (A), or by a two-way ANOVA with Sidak’s correction for multiple comparisons (J). ns = not significant, **p < 0.01, ***p < 0.001, ****p < 0.0001.

Because the boronic acid residue serves as an electrophile to generate a covalent adduct with the nucleophilic active site threonine of the β5 subunit, we hypothesized that the nucleophilic thiol of CYS could similarly react with the boronic acid of bortezomib, blocking its ability to inhibit the proteasome^20,45^. This hypothesis was further motivated by prior work with other nucleophiles such as green-tea EGCG and vitamin C, which were shown to react with bortezomib’s boronic acid and block its proteasome-inhibitory activity^46,47^. To test this possibility, we incubated bortezomib with vehicle (PBS), CYS or CYS_2_ (which does not have chemically available thiols) and performed LC-MS to look for potential fates of bortezomib, including possible conjugates between CYS and bortezomib (**Figure 4D, Extended Data Figure 8D**). We first assessed bortezomib in vehicle treated conditions, finding sodium adducts of bortezomib (m/z 407.1861) and its de-boronated degradation product (m/z 379.1741) consistent with previous observations^48,49^ (**Figure 4E-F**). These bortezomib-associated peaks are also found in samples co-incubated with CYS and CYS_2_, however the de-boronated product was substantially increased upon incubation with CYS, suggesting that CYS may mitigate bortezomib activity, in part, by catalyzing its de-boronation (**Figure 4F**). Notably, in the CYS-incubated samples only, we also identified an ion consistent with an expected covalent product between CYS and bortezomib (m/z 510.1953) (**Figure 4E-F**). An analogous de-boronated CYS-bortezomib conjugate was not detected, consistent with a hypothesized reaction between CYS and bortezomib occurring at the boronic acid (**Extended Data Figure 8D**).

Considering that incubation of bortezomib with CYS can generate covalent conjugates and promote deboronation, we next asked if these CYS interactions functionally impacted bortezomib activity. We treated cells with bortezomib that was not pre-incubated (Unmodified) or was pre-incubated with vehicle, CYS, or CYS_2_. Consistent with CYS-mediated detoxification of bortezomib, only pre-incubation of CYS diminished bortezomib efficacy (**Figure 4G**). A similar loss of efficacy from CYS pre-incubation was found with the boronic acid containing proteasome inhibitor ixazomib (**Extended Data Figure 8E**). While pre-incubation alone partially diminished the efficacy of the non-boronic acid containing carfilzomib, CYS incubation had minimal additional effect (**Figure 4H**). Together, these data suggest that free intracellular CYS can suppress cellular toxicity to boronic acid proteasome inhibitors via direct inactivation.

We next investigated how interactions between α-ketoacids, CYS, and boronic acid proteasome inhibitors impact proteasome activity. Prior studies have shown that cell death from proteasome inhibition can be rescued downstream of the proteasome by amino-acid supplementation or by glutathione-dependent redox buffering^33,40^. Given the finding that CYS can form thiazolidines with α-ketoacids, we hypothesized that α-ketoacid supplementation decreases available intracellular CYS, preventing CYS from forming conjugates with bortezomib, and leading to greater target engagement and proteasome inhibition. To examine this possibility, we measured the accumulation of total ubiquitinated protein in bortezomib-treated A549 cells treated with either the α-ketoacid pyruvate or phenylpyruvate when supplemented with 1 mM CYS. While pyruvate and phenylpyruvate did not impact levels of total ubiquitinated protein alone, supplementing cells with either α-ketoacid metabolite led to greater accumulation of ubiquitinated protein upon low dose bortezomib treatment that was reversed by CYS co-treatment (**Figure 4I**). We then directly measured the chymotrypsin-like activity of the proteasome in cells treated under these conditions. Consistently, both pyruvate and phenylpyruvate enhanced the inhibition of chymotrypsin-like activity upon bortezomib treatment, which was mitigated by co-treatment with CYS (**Figure 4J**). Importantly, CYS alone did not affect chymotrypsin-like activity of cells in the absence of bortezomib, nor did CYS rescue the proteasome activity of carfilzomib-treated cells (**Extended Data Figure 8F-G**). These findings suggest that both α-ketoacid and CYS availability modulate cell sensitivity to boronic acid proteasome inhibitors by influencing proteasome target engagement. Altogether, we propose a model where α-ketoacids, such as pyruvate, influence cell sensitivity to boronic acid proteasome inhibitors by forming thiazolidines that alter intracellular CYS availability. Because CYS can form conjugates with bortezomib and promote deboronation, CYS suppresses boronic acid proteasome inhibitor activity. Thus, the generation of thiazolidines upon α-ketoacid treatment lowers effective CYS concentrations, preventing drug detoxification and increasing proteasome inhibition at a given concentration of boronic acid proteosome inhibitor (**Figure 5**).

**Figure 5.**
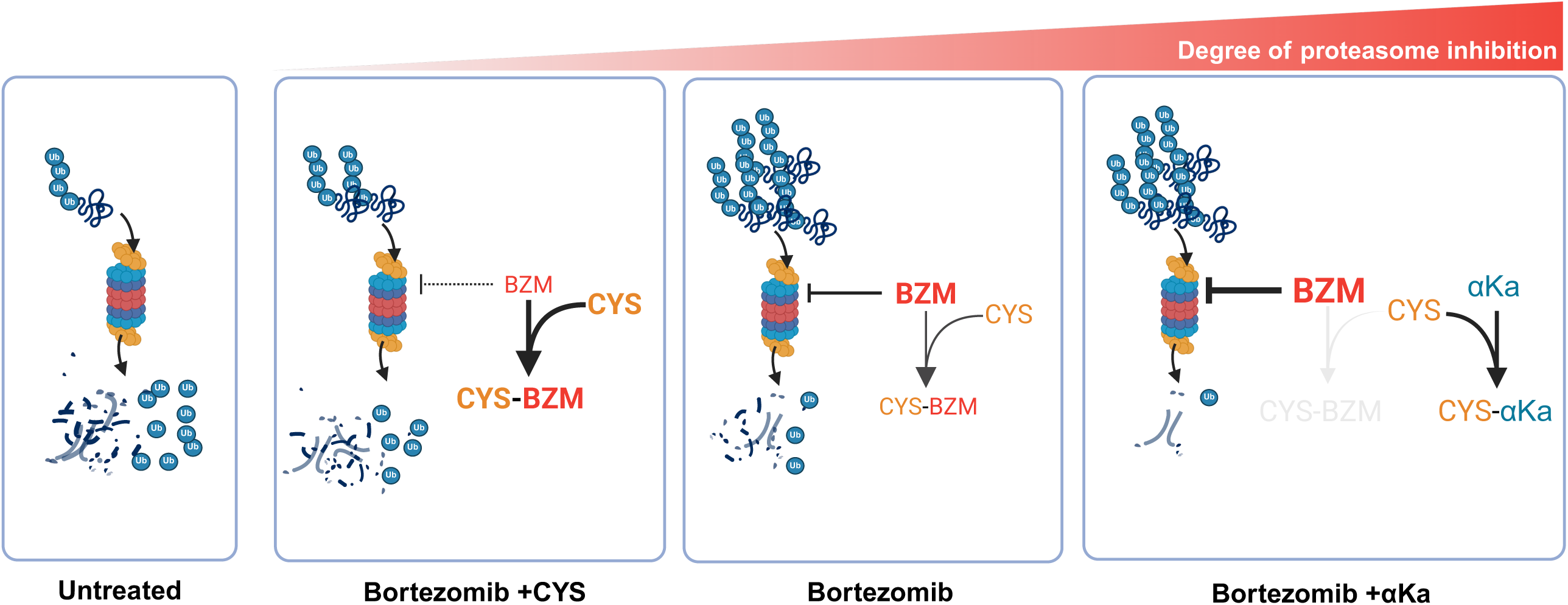
Proteasome inhibition by bortezomib is governed by the availability of CYS and α-ketoacids. Schematic depicting the proposed mechanism of the interactions between CYS, α-ketoacids, and boronic acid-containing proteasome inhibitors to impact the degree of proteasome activity.

## Discussion

In this study, we reveal that α-ketoacids, including the metabolite pyruvate, increase cell sensitivity to clinically approved boronic acid proteosome inhibitors such as bortezomib. Additionally, we elucidate a mechanism by which CYS confers cell resistance to boronic acid proteasome inhibitors through direct chemical antagonism to suppress proteasome inhibition function. Our results suggest that CYS can detoxify bortezomib both by catalyzing de-boronification of the drug, as well as forming a covalent conjugate. These two mechanisms underscore the importance of limiting boronic acid proteasome inhibitors from reaction with CYS. By investigating the chemical features and interactions between CYS, boronic acid proteasome inhibitors, and α-ketoacids, we find that α-ketoacids react with CYS to decrease free intracellular CYS availability, functionally increasing the availability of boronic acid proteasome inhibitors to engage their target.

CYS plays a role in modulating therapeutic responses to many drugs. Several cancer contexts, including those with constitutive activation of the stress response transcription factors ATF4 or NRF2, typically have increased expression of SLC7A11, the rate-limiting component of xCT and its associated CYS_2_ uptake^42,50–52^. Interestingly, cancers with activation of ATF4 or NRF2 have been associated with drug resistance, suggesting that CYS dysregulation may be a recurring contributor to drug resistance phenotypes. In the case of plasma cell dyscrasias SLC7A11 expression increases with disease progression^53^, indicating that increased CYS may contribute to bortezomib resistance in this context. If so, these findings suggest two potentially actionable considerations: 1) Selection of PROTEASOME INHIBITOR drugs based on the likelihood for CYS-mediated boronic acid PROTEASOME INHIBITOR resistance, 2) Co-treatment of α-ketoacids as a potential strategy to enhance boronic acid PROTEASOME INHIBITOR efficacy. Indeed, in the latter case recent work found that systemic pyruvate administration increased bortezomib efficacy in mice^15^, highlighting that treatment with α-ketoacids may be a general strategy to improve proteosome inhibitor drug efficacy.

Boronic acids, due to their ability to form reversible covalent bonds, reactive versatility, and low toxicity, are important functional groups for small molecule therapeutics^54^. Beyond bortezomib and ixazomib, several other clinically relevant agents also possess boronic acids, including the antibacterial vaborbactam and the antifungal benzoxaborole. Our work suggests that microbial levels of CYS and α-ketoacids may similarly impact the efficacy of these boronic acid-containing compounds. Furthermore, analogous processes may also occur with other, non-boronic acid containing electrophilic drugs across a wide swath of therapeutic contexts. Overall, this work highlights how complex chemical interactions between metabolites and therapeutic compounds can impact drug efficacy.

## Methods

### Quantitative High-Throughput Small Molecule Screening

Drug screening was performed as described in Griner et al. PNAS 2014 with the following changes^55^. 2480 compounds from the MIPEv5.0 library were tested at 11 concentrations in a 1:3 dilution series starting at 40 µM^35^. DMSO was used as a negative control and 2 µM bortezomib was used as a positive control for cytotoxicity. On day 0, A549 cells were seeded in 1536-well plates white polystyrene tissue culture-treated plates (Corning) at a density of 500 cells/well, using a Multidrop Combi dispenser (ThermoFisher). Seeding volume was of 5 µL/well of DMEM without pyruvate supplemented with 10% dialyzed FBS (VWR). Cells were then incubated overnight in a 5% CO2 incubator to enable reattachment to the plate On day 1, pyruvate was added to each well to a final concentration of 0 or 5 mM and library compounds were added to each well at the desired concentration via a 1536 pin-tool. Plates were incubated for 48 hours at standard incubator conditions covered by a stainless steel gasketed lid to prevent evaporation. For assessment of cell viability, 3 µL of CellTiter Glo (Promega) reagent was added to each well, plates were incubated for 15 minutes at room temperature. Luminescence signals were then measured on a Viewlux imager (PerkinElmer) with a 2 s exposure time per plate. Relative luminescence units (RLUs) for each well were growth-rate corrected using the following equation: 50*2^log(RLUdrug/RLUt0)/log(RLUDMSO/RLUt0)^ where RLU_DMSO_ represents the median DMSO signal at the final timepoint, and RLU_t0_ represents the median signal across wells at the timepoint just before pyruvate and drug addition. In this equation, a corrected growth rate of 100 corresponds to equivalent proliferation in drug compared to DMSO, 50 indicates zero proliferation, and 0 indicates complete loss of viability. For analysis of sensitivity to pyruvate concentration, only drugs with at least a 20% drop in growth rate at the highest drug concentration were considered, and area under the curve (AUC) was calculated by summing corrected growth rates for the highest 8 concentrations and subtracting the corrected growth rates for the lowest 4 concentrations (to remove technical artifacts for some of the lower concentrations).

### Cell Culture

Cell lines were acquired from ATCC (A549), Leibniz Institute DSMZ (L-363, AMO-1), as a gift from Dr. Supriya Saha (CCLP1), or as a gift from Dr. Sita Kugel (SUIT2). Cell identities were confirmed using STR profiling and cells were regularly tested to be free of mycoplasma contamination (MycoProbe, R&D Systems). Cells were sustained in Dulbecco’s Modified Eagle Medium (DMEM) supplemented with 3.7 g/L sodium bicarbonate (Sigma, S6297), 10% heat-inactivated fetal bovine serum (FBS) (Cytiva HyClone SH3039603HI), and 1% penicillin/streptomycin solution (P/S) (Sigma, P4333). Cells were incubated in a humidified incubator at 37 C and 5% CO_2_.

### Media Conditions and Treatments

Cells were seeded into 96-well or 6-well plates depending on the experiment. The following day, wells were aspirated and changed to DMEM assay media containing dialyzed FBS (Sigma, F0392) and various treatments. Treatments included varying concentrations of pyruvate (Sigma Aldrich, P8574), phenylpyruvic acid (Sigma Aldrich, 286958), alpha-ketoglutaric acid (Sigma Aldrich, K2010), 2-ketobutyric acid (Millipore Sigma, K401), mesoxalate (Medchem Express HY-W540188A), L-lactate (Sigma Aldrich, L7022), 500 µM supplemental cystine (high cystine) (Sigma Aldrich, C6727), cysteine (Sigma Aldrich, 30089). Drugs used include bortezomib (Medchem Express, HY-10227), Ixazomib (Medchem Express, HY-10453), carfilzomib (Medchem Express, HY-10455), MG-132 (Selleck Chemicals, S2619), MG-262 (ApexBio, A8179), sodium dichloroacetate (Sigma Aldrich, 247795), and UK-5099 (Cayman, 16980).

### Cell Growth Assays

For growth assays using A549, CCLP1, and SUIT2, cells were seeded at 1,000-2,000 cells per well in standard media conditions in 96-well plates (Fisher Scientific, FB012931). After overnight attachment (day 0), an untreated 96-well plate was fixed with 10% TCA (Fisher, A11156.0B) to serve as a control. After 4 days, all treated plates were similarly fixed with 10% TCA. Plates were left in 10% TCA for 1-4h, then rinsed 6x in DI water and 75 µL of 0.057% sulforhodamine B (SRB) (Sigma, 230162) in 1% acetic acid was added to each well for 0.5-4h. SRB was then decanted out of the plates and rinsed 3x times with 150 µL of 1% acetic acid per well (Sigma, AX0074-6). SRB was then redissolved in 150 µL of 10 mM Tris (pH 10.5). Plates were read on a Tecan plate reader to measure absorbance at 510 nm. Growth rate was determined by the following equation:

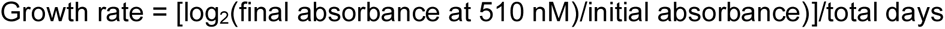

For growth assays using suspension cells L-363, AMO-1, and K562, cells were plated at 40,000 cells per well in 2 mL of standard media conditions in 12-well plates. The next day, treatment media was added to each well to obtain final treatment concentrations. Untreated cells on day 0 and treated cells after 72 hours of treatment were counted using a Multisizer 3 Coulter Counter (Beckman Coulter). Growth rate was determined by the following equation:

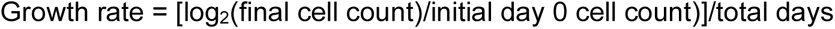

### Metabolite Extractions

Cells were kept on ice throughout the whole extraction process. First, spent media was aspirated, then the wells were washed once with blood bank saline, which was then aspirated. Then, 500 µL of extraction solvent was added to each well, wells were scraped using the back of a p1000 pipette tip, and the whole volume was transferred into a microcentrifuge tube and placed on ice. Depending on the goal of the experiment, the extraction solvent was varied. For standard extractions, 80% methanol (80:20 methanol:water) with or without valine D8 loading standard was used. For detection of thiol metabolites, a solution of 80% methanol with 20% 10 mM ammonium formate (pH 7) in HPLC grade water with 2.5 mM N-ethylmaleimide (final concentration 2 mM ammonium formate, 0.5 mM NEM). All samples were centrifuged at 17,000 g for 5 minutes at 4C, and 350 µl of supernatant was transferred into a fresh microcentrifuge tube and dried on a Centrivap vacuum concentrator (Labconco, 10269602). Replicate wells for each extracted condition were also counted on a Coulter Counter to determine the total average cell volume for each condition. For analysis, cell extracts were resuspended in 80% MeOH at a concentration of 28 µL solvent per 1 µL cell volume, vortexed at 4C for 10 minutes, and centrifuged at 17,000 g for 10 min at 4C. 20 µL of metabolite extract was transferred into an LCMS vial and stored at -20C until analysis.

### Generation of CYS-drug conjugates by combining CYS with proteasome inhibitors

bortezomib was solubilized to 200 mM in DMSO and carfilzomib to 150 mM in DMSO. L-cysteine (CYS) was prepared to 15 or 20 mM in PBS that had been degassed for >1hr. Reactions were prepared in duplicate by mixing 100 µL of CYS with 1 µL of drug for a final molar ratio of 10:1 Cys:bortezomib. Controls were included by adding 100 µL of PBS or CYS_2_ (10:1 molar ratio), as indicated. Tubes were vortexed, gently centrifuged to bring contents to the bottom of the tube, and incubated at RT overnight. After incubation, 400 µL 100% MeOH was added to each tube and vortexed for a final concentration of 80% MeOH. Tubes were centrifuged at max speed for 5 mins and 400 µL was transferred to a fresh tube and dried down. Once dried, 1600 µL of 80% MeOH was added to each tube, vortexed at 4C for 10 min, and 20 µL was transferred to an LCMS vial for analysis.

### LC-MS

Untargeted metabolite quantitation of resolubilized metabolite extracts from cells was performed using a Q Exactive HF-X Hybrid Quadrupole-Orbitrap Mass Spectrometer equipped with an Ion Max AP source and H-ESI II probe, coupled to a Vanquish Flex Binary UHPLC system (Thermo Scientific).

### Untargeted Metabolomics

Metabolite quantitation of resolubilized metabolite extracts was performed using a Q Exactive HF- X Hybrid Quadrupole-Orbitrap Mass Spectrometer equipped with an Ion Max API source and H- ESI II probe, coupled to a Vanquish Flex Binary UHPLC system (Thermo Scientific). Mass calibrations were completed at a minimum of every 5 days in both the positive and negative polarity modes using LTQ Velos ESI Calibration Solution (Pierce). Metabolites were chromatographically separated by injecting a sample volume of 1 μL into a SeQuant ZIC-pHILIC Polymeric column (2.1x150 mm 5 mM, EMD Millipore). The flow rate was set to 150 µL/min, autosampler temperature set to 10 °C, and column temperature set to 30 °C. Mobile Phase A consisted of 20 mM ammonium carbonate and 0.1% (v/v) ammonium hydroxide, and Mobile Phase B consisted of 100% acetonitrile. The sample was gradient eluted (%B) from the column as follows: 0–20 min.: linear gradient from 85% to 20 % B; 20–24 min.: hold at 20% B; 24–24.5 min.: linear gradient from 20% to 85% B; 24.5 min.-end: hold at 85% B until equilibrated with ten column volumes. Mobile Phase was directed into the ion source with the following parameters: sheath gas = 45, auxiliary gas = 15, sweep gas = 2, spray voltage = 2.9 kV in the negative mode or 3.5 kV in the positive mode, capillary temperature = 300 °C, RF level = 40%, auxiliary gas heater temperature = 325 °C. Mass detection was conducted with a resolution of 240,000 in full scan mode, with an AGC target of 3,000,000 and maximum injection time of 250 msec. Metabolites were detected over a mass range of 70–1050 m/z. Quantitation of all metabolites was performed using Tracefinder 4.1 (Thermo Scientific) referencing an in-house metabolite standards library using ≤5 ppm mass error.

### Targeted metabolomics

Quantification of resolubilized metabolite extracts was performed on an Agilent 6495D Triple-Quadrupole mass spectrometer equipped with an Agilent JetStream Heated ESI source, coupled to an Agilent 1290 Infinity II UHPLC system. A check tune was performed using the Agilent ESI-L low concentration tuning mix to assess the instrument performance prior to data collection. Polar metabolite samples were chromatographically separated by loading a 5 µL injection volume onto one of two paired (K’ value matched) Poroshell 120 HILIC-Z columns (2.1x150mm, 2.7 micron, Agilent Technologies) running in an alternating column regeneration method. The flow rate was 600uL/min, column temperature was set to 25C, and the multisampler temperature was set to 4C. Mobile phase “A” consisted of 0.1% formic acid (v/v in H2O) with 10mM ammonium formate, and mobile phase “B” was acetonitrile with formic acid at 0.1% (v/v). The sample was gradient eluted from the column as follows: 0-0.14 min, initial hold at 95% “B”; 0.14-2.29 min, 95% to 40% “B”; 2.29-4.0 min, hold at 40% “B”. At the end of the method, the valve in the column oven would switch over to the other HILIC-Z column and the gradient would run on column 2, while column 1 was regenerated at 40% to 95% “B” from 0-0.56min, followed by an increase in flow-rate to 1200 µL/min and held at 95% “B” until 10 column volumes were pumped through the column. Samples were analyzed on the mass spectrometer using the following parameters: gas flow = 13.0L/min, sheath gas = 11L/min, nebulizer = 35psi, gas temperature = 200C, sheath gas temperature = 250C, capillary voltage = 3000V, nozzle voltage = 1500V and CAV voltage of 5V. Metabolites were targeted in the MRM mode with a dwell time of 100ms at unit resolution; compound collision energies (CEs) were previously optimized in the automated mode of MassHunter’s built in optimizer module (ver. 12.1). Quantification of all metabolites was performed using MassHunter’s Quantitative Analysis module referencing an in-house metabolite standards library and an isotopically labelled standard.

### Immunoblotting

Cells were washed with ice cold 1X PBS and lysed in ice cold RIPA buffer containing Halt Protease and Phosphatase Inhibitor Cocktail (Thermo Scientific 78442). Lysates were clarified by rocking samples for thirty minutes at 4°C and then spun at maximum speed for ten minutes for collection of supernatant. Protein concentration was calculated using the BCA Protein Assay (Pierce, 23225) with BSA as a standard. Lysates were resolved by SDS-PAGE using NuPAGE 4-12% Bis-Tris Protein Gels and run at 100V. Proteins were transferred onto nitrocellulose membranes via wet transfer at 100V. Membranes were blocked in 5% BSA in TBST before incubating membranes with primary antibodies at 4°C overnight (Vinculin [1:1000] Cell Signaling Technology 13901S; Ubiquitin [1:1000] Cell Signaling Technology 3933S; GAPDH) [1:3000] Cell Signaling Technology D16H11; Ubiquitin (P4D1) [1:500] Santa Cruz Biotechnology sc-8017). The next day, membranes were washed three times with TBST at room temperature and then incubated in secondary antibodies for one hour at room temperature. The secondary antibodies used were anti-rabbit IgG horseradish peroxidase-linked antibody (1:4000 dilution; Cell Signaling Technologies, 7074S, anti-mouse IgG horseradish peroxidase-linked antibody (1:4000, 7076S)), IRDYE 800CW goat anti-mouse IGG secondary antibody (1:10000 dilution; LICORbio, 926-32210), or IRDYE 680CW goat anti-rabbit IGG secondary antibody (1:10000 dilution; LICORbio, 926-68071)

### Cell-Based Proteasome Activity Assays

Proteasome activity was measured using the Cell-Based Chymotrypsin-like Proteasome-Glo Assay (Promega G8660) per manufacturer’s protocol. Briefly, cells were trypsinized and washed three times with 1X PBS before plated. 4,000 cells were plated in a white-walled 96 well plate and allowed to settle in 250 μL of media containing 10% dialyzed FBS. The next day, cells were washed twice in 200 μL of treatment media before 100 μL of treatment was added for the indicated amount of time. One hour before initiating the assay, the cell plate was taken out of the incubator and equilibrated at room temperature. 100 μL of luciferin detection reagent with chymotrypsin-like substrate was added to each well with the exception of substrate-free controls, incubated in the dark for ten minutes, and then placed in a plate reader (Tecan Infinite M200Pro) to measure luminescence.

### Generation of UbG76V-GFP Reporter Cells

AMO-1 cells expressing UbG76V-GFP were generated via lentiviral infection. pLJM1-UbG76V-GFP was constructed using pLJM1-EGFP (Addgene 19319) as a backbone. pLJM1-EGFP was digested using AgeI and EcoRI and gel purified to remove the EGFP sequence. UbG76V-GFP cDNA was PCR amplified using Phusion High-Fidelity DNA polymerase (NEB M0530) from the parental UbG76V-GFP plasmid (Addgene 11941) with the following oligonucleotide primers:

UbG76V-GFP F: tttagtgaaccgtcagatccgctagcgctaccggtcgccaccatgcagatcttcgtgaagac UbG76V-GFP R:
Gatgaatactgccatttgtctcgaggtcgagaattcgaagcttgagctcgagatctgagtccggattacttgtacagctcgtcc\

Digested pLJM1 backbone and amplified UbG76V-GFP were assembled using Gibson Assembly, and the resulting plasmid was transformed and validated by Sanger sequencing. Lentiviral production was done by transfecting constructs into LentiX293T cells with Mirus Transit293T (Mirus Bio MIR2700) following manufacturer’s protocol. AMO-1 cells were infected and selected with puromycin (Sigma Aldrich P7255). UbG76V-GFP expression was validated by treating infected cells and parental AMO-1 cells with bortezomib and performing flow cytometry to observe selective GFP accumulation in infected cells. All experiments were conducted on a polyclonal cell population.

### Flow Cytometry Analysis for UbG76V-GFP Accumulation

AMO-1 cells expressing UbG76V-GFP were plated at a density of 10,000 cells per well in a 96-well plate in 100 μl of treatment media. Cells were treated with either 0 or 4 nM bortezomib or carfilzomib for 0, 8, and 24 hours. Treated cells were then spun down at 1000xg for 3 minutes, resuspended in 100 μl of 1X PBS, spun down again at 1000xg for 3 minutes, followed by staining with 1 μg/mL DAPI (Thermo Fisher D1306) in Fluorobrite DMEM (Thermo Fisher A1896701). Samples were immediately analyzed for viability and GFP accumulation. Flow cytometry analysis was performed on the BD FACSCanto machine. Data were analyzed by FlowJo (v10.8.0). Uninfected parental AMO-1 cells were used as a negative control.

### Purified 20S Proteasome Activity Assays

Chymotrypsin-like proteasome activity of purified human 20S proteasome (Carrier Free; R&D Systems, E-360) was measured using the Proteasome-Glo Assay system following the manufacturer’s protocol. Briefly for each sample, 0.5 μg of protein was resuspended in 50 μl 10 mM HEPES with treatment as indicated. 0.5 μg was chosen after doing a standard curve to measure the assay limit of detection. Samples were incubated at 37C for 1 hour, then returned to room temperature before 50 μL of room temperature luciferin detection reagent with chymotrypsin-like substrate was added with the exception of substrate-free controls. Samples were then incubated in the dark for ten minutes and placed in a plate reader (Tecan Infinite M200Pro) to measure luminescence.

### Statistics

Statistical tests used across experimental groups are annotated in each figure legend were conducted in Graphpad Prism 10, with exact test results provided in the supplementary files. For growth curves, statistical significance was assessed by fitting a nonlinear variable slope (four parameters) regression model with an extra sum-of-squares F test and results are provided in the supplementary files. Sample sizes were not pre-determined but were based on observed variance in standard measurements (LC-MS experiments, growth assays). Data distribution was assumed to be normal, but this was not formally tested. When possible, samples groups were randomized in the order of analysis to distribute systemic errors. Data collection and analysis were not performed blind to the conditions of the experiments. All experiments have been repeated at least once with qualitatively similar results. All measurements shown are from distinct samples, with datapoints representing technical replicates from parallel conditions on the same experiment, unless stated otherwise.

### Contributions

JAB, SMC, MK, ZL, LBS, MVH conceived and designed experiments. JAB, SMC, MK, LGR, SK, ZL, KJC, SDB, RC, GEO performed the experiments. MC, DH, BTD, and CJT performed and analyzed the small molecule screen. JAB, SMC, MK, SK, ZL, SDB, LBS, MVH analyzed the data. JAB, SMC, MK, LBS, MVH contributed materials/analysis tools. JAB, SMC, LBS wrote the paper.

## Acknowledgements

We thank the Vander Heiden and Sullivan laboratories for scientific discussion. This research was supported by the Proteomics & Metabolomics Shared Resource, which is supported by a National Cancer Institute Cancer Center Support Grant for the Fred Hutch/University of Washington/Seattle Children’s Cancer Consortium (NCI: P30CA015704) and by the Koch Institute Flow Cytometry Core Facility (NCI: P30CA014051). L.B.S. acknowledges support from the NIGMS (R35GM147118). J.A.B acknowledges support from NIGMS T32GM007270 and a Graduate Student Research Fellowship from the National Science Foundation (DGE-2140004). S.M.C acknowledges support from the Ruth L. Kirschstein National Research Service F30 Award (F30CA268633) and the Harvard/MIT MSTP (T32GM007753, T32GM007287). M.K. acknowledges support from the Levinson Emerging Scholars Award. Z.L. acknowledges support from the National Institutes of Health (T32GM007287). B.T.D. also acknowledges support from MSTP T32GM007753) M.G.V.H. acknowledges support from the Ludwig Center at MIT, the MIT Center for Precision Cancer Medicine, and the NCI (R35CA242379, R01CA259253, P30CA014051). This research was supported in part by the Intramural Research Program of the National Institutes of Health (NIH). The contributions of the NIH authors are considered Works of the United States Government. The findings and conclusions presented in this paper are those of the authors and do not necessarily reflect the views of the NIH or the U.S. Department of Health and Human Services. The funders had no role in study design, data collection and analysis, decision to publish or preparation of the manuscript. Four figure panels were created in Biorender by J.A.B. (Figures 1A, 2A, 4D, 5), licensed under CC BY 4.0: https://biorender.com/8pt1ayf. All chemical structures were created in ChemDraw 22.2.0 by J.A.B. We thank Brian Milless and Jared Mayers for continuous technical support with LC-MS experiments. We additionally thank Julia Crainic for assistance in data visualization.

## Data Availability

All data supporting the findings of this study are available within the paper and its source data files. Unprocessed western blot data, data used to generate all figures, and statistical test results are included in the source data files.

---

**Extended Data Figure 1.**
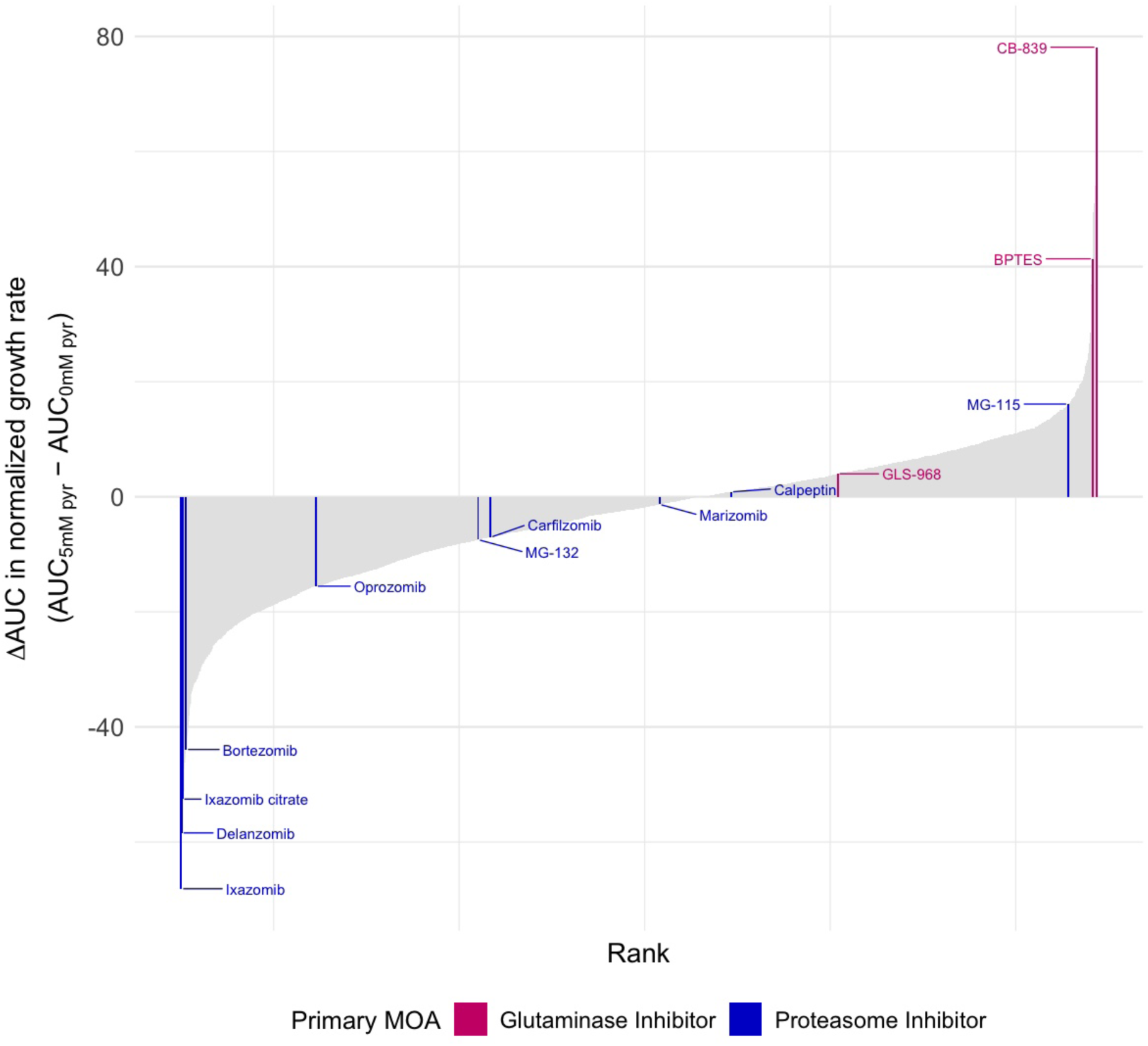
Results from Quantitative High-Throughput Small Molecule Screening. **(A)** Small molecules of interest are color-coded by the different mechanisms of actions, as indicated. Pyruvate increased cell resistance to small molecules where *Δ*AUC is positive. Pyruvate increased cell sensitivity to small molecules where *Δ*AUC is negative.

**Extended Data Figure 2.**
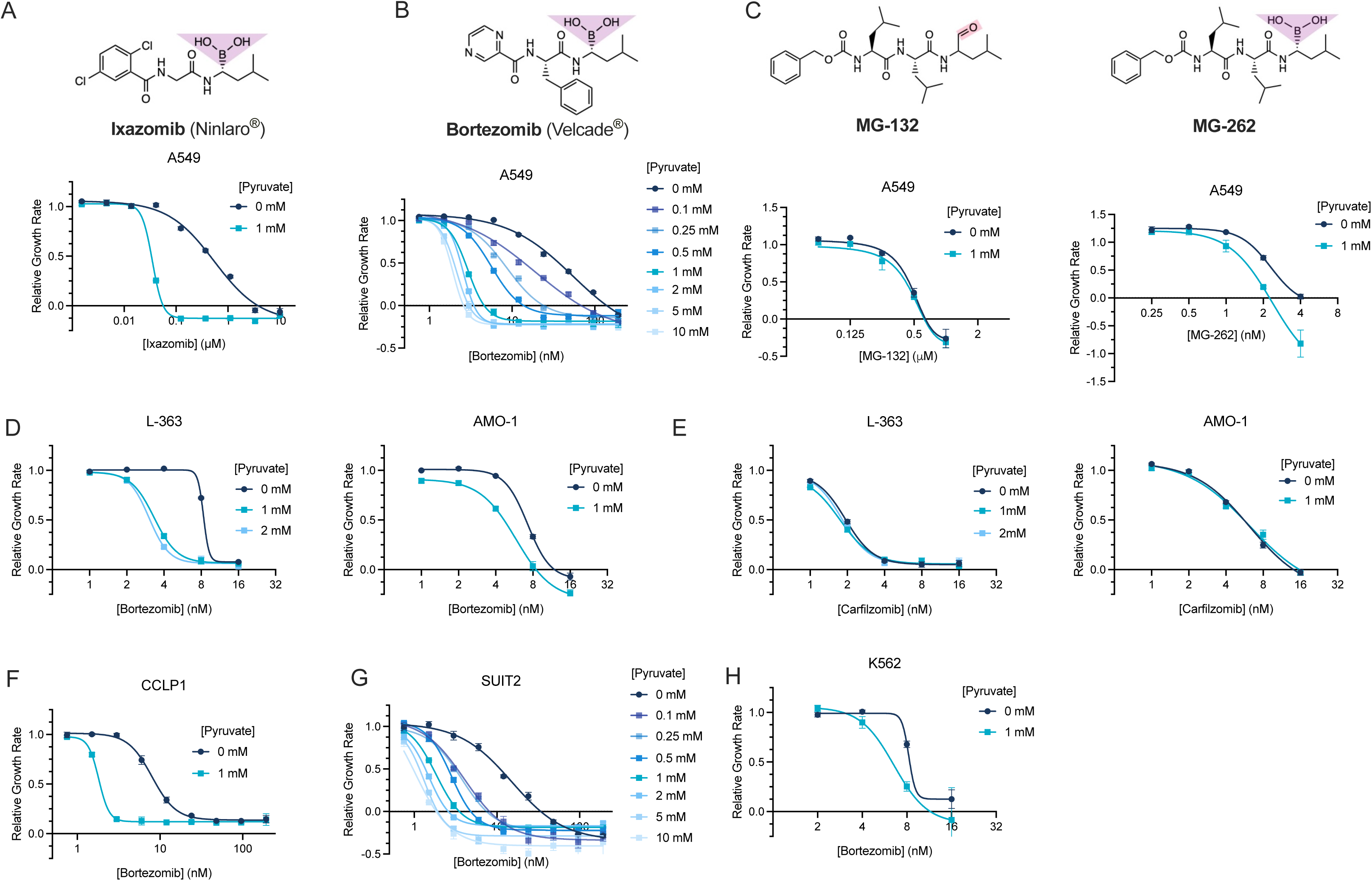
Pyruvate increases cell sensitivity to bortezomib in multiple cancer cell types in a dose-dependent manner. **(A)** Above: chemical structure of ixazomib; Below: relative growth rate of A549 cells treated with or without 1 mM pyruvate and cotreated with varying doses of ixazomib for 96 hours. Growth rates are relative to the untreated conditions, n=6 per condition. **(B)** Above: chemical structure of bortezomib; Below: relative growth rate of A549 cells treated with pyruvate from 0 mM to 10 mM and cotreated with varying doses of bortezomib for 96 hours. Growth rates are relative to the untreated conditions, n=3 per condition. **(C)** Above: structures of MG-132 (left) and MG-262 (right) with the functional groups highlighted by chemical class as denoted in Figure 1B. Below: relative growth rate of A549 cells treated with or without 1 mM pyruvate and cotreated with increasing doses of MG-132 (left) or MG-262 (right) for 72 hours, Growth rates are relative to the untreated conditions, n=2 per condition. **(D)** Relative growth rate of L-363 or AMO-1 cells treated with the indicated concentrations of pyruvate and cotreated with increasing doses of bortezomib for 72 hours. Growth rates are relative to untreated conditions, n=3 per condition. **(E)** Relative growth rate of L-363 or AMO-1 cells treated with the indicated concentrations of pyruvate and cotreated with increasing doses of carfilzomib for 72 hours. Growth rates are relative to the untreated conditions, n=3 per condition. **(F)** Relative growth rate of CCLP1 cells treated with or without 1 mM pyruvate and cotreated with increasing doses of bortezomib for 96 hours. Growth rates are relative to the untreated condition, n=6 replicate per condition. **(G)** Relative growth rate of SUIT2 cells treated with indicated concentrations of pyruvate and increasing doses of bortezomib for 96 hours. Growth rates are relative to the untreated condition, n=3 per condition. **(H)** Relative growth rate of K562 cells treated with indicated concentrations of pyruvate and increasing doses of bortezomib for 72 hours. Growth rates are relative to the untreated condition, n=3 per condition. For all panels values represent means, error bars are SEM. Relative growth rate curves were fitted with a nonlinear variable slope (four parameters) regression model.

**Extended Data Figure 3.**
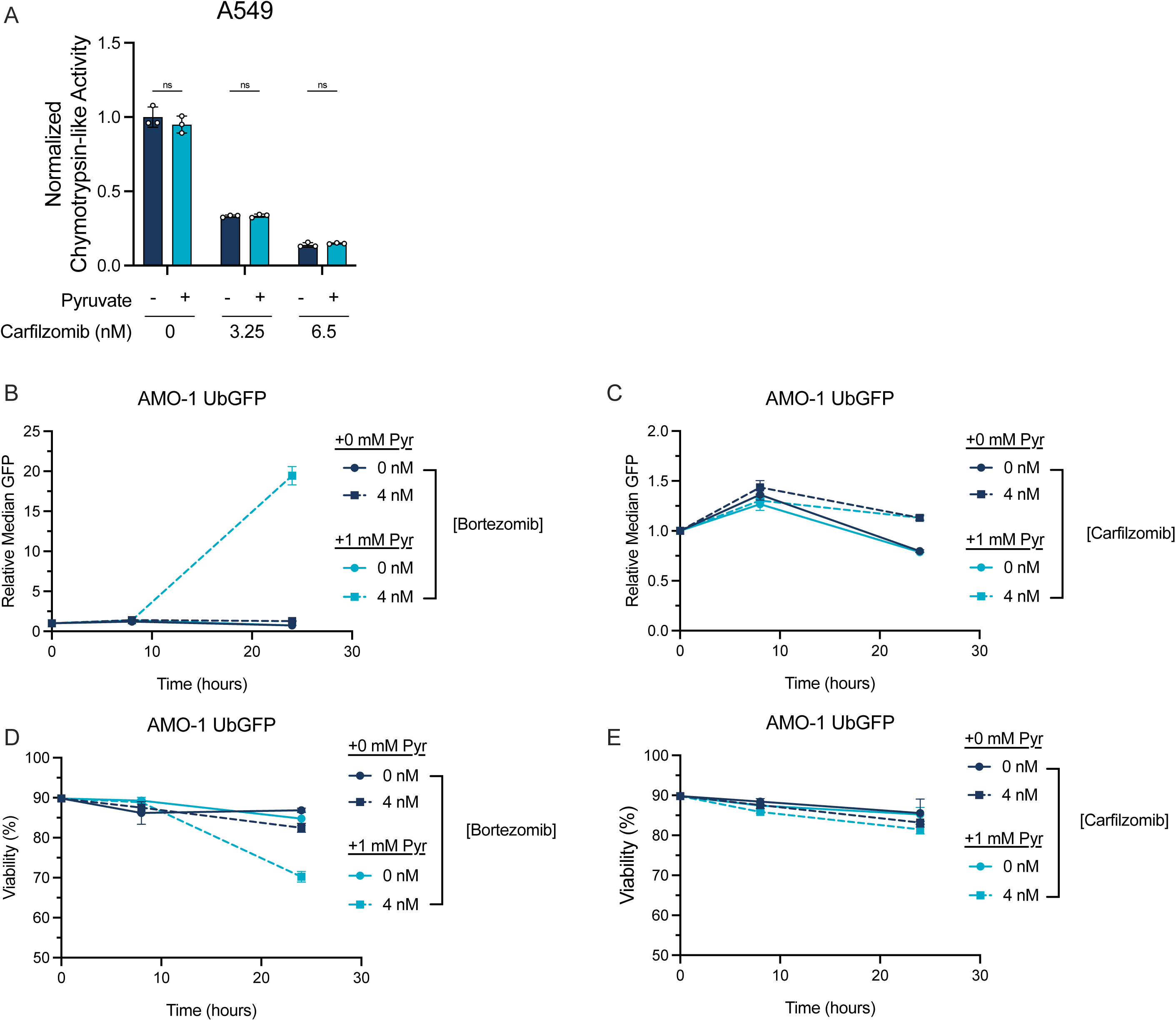
Pyruvate supplementation suppresses ubiquitin-proteasome dependent proteolysis in the presence of bortezomib. **(A)** Cell-based proteasome activity assay for chymotrypsin-like activity in A549 cells treated with or without 1 mM pyruvate or carfilzomib for 6 hours, as indicated. Values were normalized to untreated conditions for each treatment condition and represent means, n=3 per condition. **(B,C)** Median GFP signal measured via flow cytometry in AMO-1 cells expressing UbG76V-GFP after 0, 8, and 24 hours of co-treatment with pyruvate and bortezomib (B) or carfilzomib (C), as indicated. GFP signal is normalized to percent viability, n=3 per condition. **(D,E)** Percent viability measured via DAPI staining in AMO-1 cells expressing UbG76V-GFP after 0, 8, and 24 hours of co-treatment with pyruvate and bortezomib (D) or carfilzomib (E), as indicated, n=3 per condition. For all panels values represent means, error bars are SEM. Statistical significance was assessed a two-way ANOVA with Sidak’s multiple comparison test (A). ns = not significant.

**Extended Data Figure 4.**
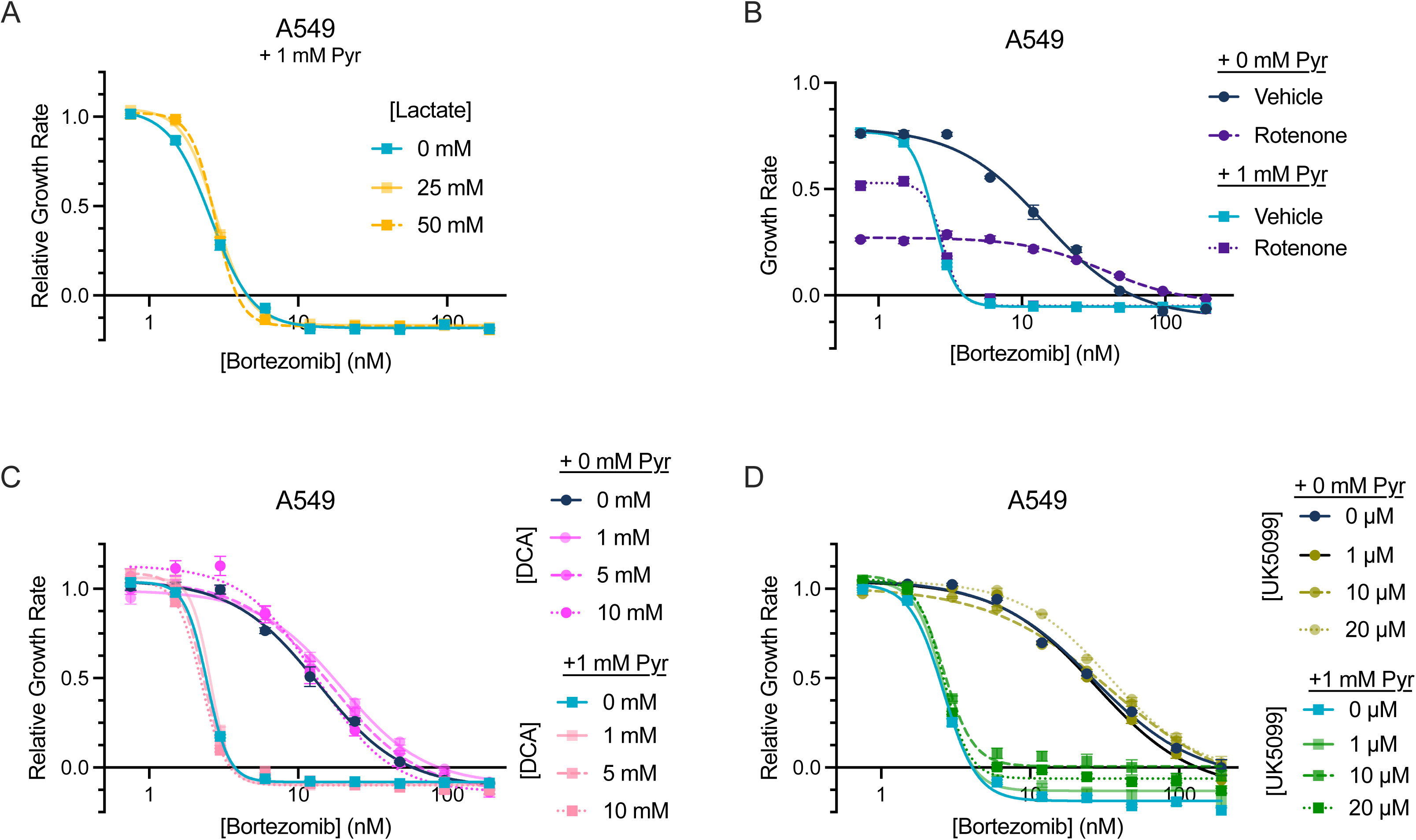
Pyruvate increases cell sensitivity to bortezomib independent of changes to mitochondrial metabolism. **(A)** Relative growth rate of A549 cells treated with or without 25 mM or 50 mM lactate and cotreated with increasing doses of bortezomib in cell culture media containing 1 mM pyruvate. Growth rates are relative to the untreated condition, n=3 per condition. **(B)** Growth rate of A549 cells treated with or without 50 nM rotenone or 1 mM pyruvate cotreated with increasing doses of bortezomib for 96 hours, n=6 per condition. **(C)** Relative growth rate of A549 cells treated with or without indicated concentrations of dichloroacetate (DCA) and 1 mM pyruvate cotreated with increasing doses of bortezomib. Growth rates are relative to the untreated condition, n=6 per condition. **(D)** Relative growth rate of A549 cells treated with or without indicated concentrations of UK5099 and 1 mM pyruvate cotreated with increasing doses of bortezomib for 96 hours. Growth rates are relative to the untreated condition, n=6 per condition. For all panels values represent means, error bars are SEM. Growth rate curves were fitted with a nonlinear variable slope (four parameters) regression model.

**Extended Data Figure 5.**
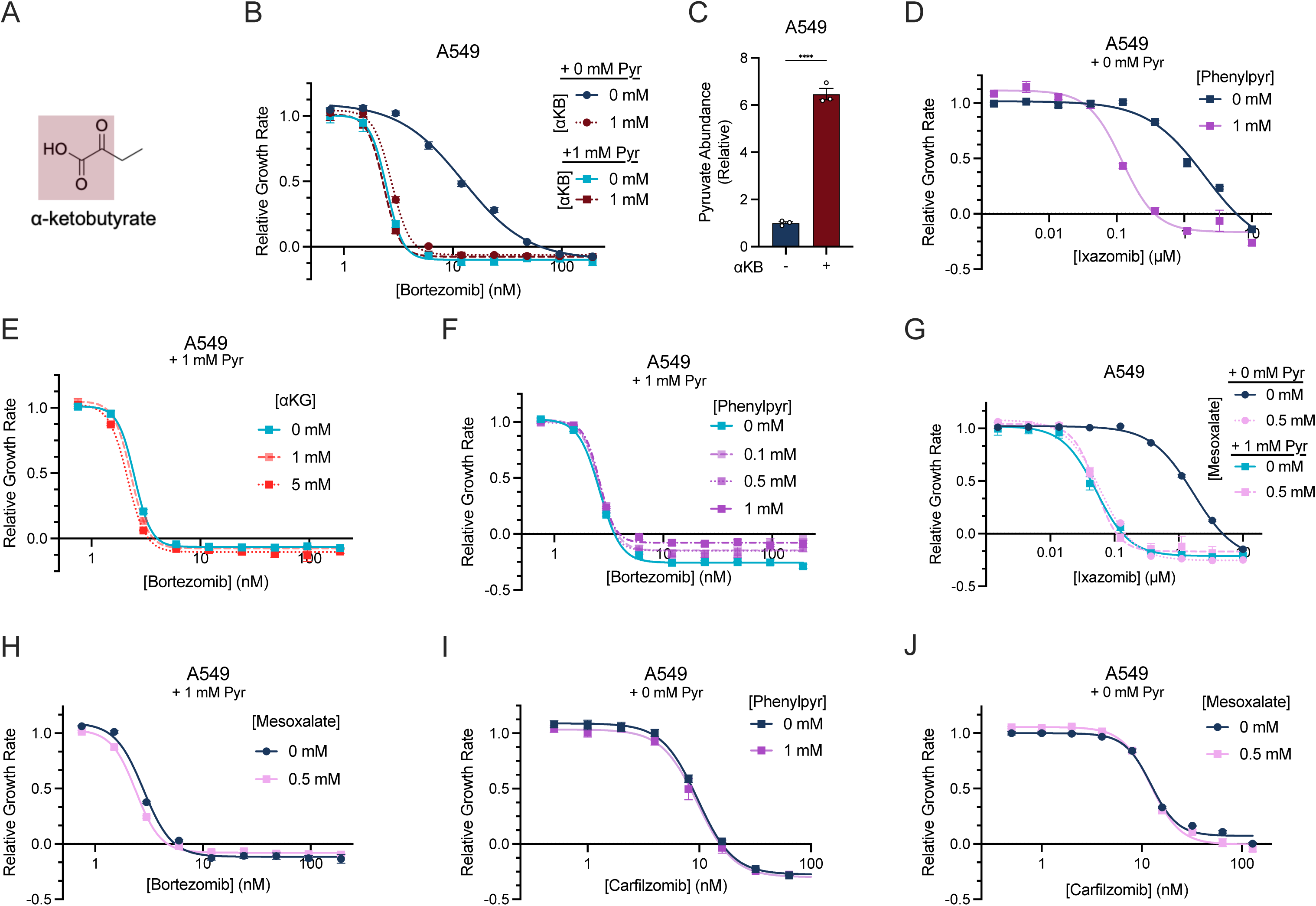
α-ketoacids increase cell sensitivity to boronic acid containing proteasome inhibitors bortezomib and ixazomib, but not the epoxyketone proteasome inhibitor carfilzomib. **(A)** Chemical structure of the α-ketoacid α-ketobutyrate (αKB) with the α-ketoacid highlighted in maroon. (B) Relative growth rate of A549 cells treated with or without 1 mM αKB and cotreated with increasing doses of bortezomib for 96 hours in media with or without 1 mM pyruvate. Growth rates are relative to the untreated condition, n=6 per condition. **(C)** Relative abundance of intracellular pyruvate as measured by LC-MS from A549 cells treated with 1 mM αKB. Abundances are relative ion counts to untreated cells, n=3. **(D)** Relative growth rate of A549 cells treated with or without 1 mM phenylpyruvate and cotreated with varying doses of ixazomib for 96 hours. Growth rates are relative to the untreated condition, n=6 condition. **(E)** Relative growth rate of A549 cells treated with or without 1 mM or 5 mM αKG and cotreated with increasing doses of bortezomib in cell culture media containing 1 mM pyruvate for 96 hours. Growth rates are relative to the untreated condition, n=6 per condition. **(F)** Relative growth rate of A549 cells treated with or without 0.1 mM, 0.5 mM or 1 mM phenylpyruvate and cotreated with increasing doses of bortezomib in cell culture media containing 1 mM pyruvate for 96 hours. Growth rates are relative to the untreated condition, n=6 per condition. **(G)** Relative growth rate of A549 cells treated with or without 0.5 mM mesoxalate and cotreated with increasing doses of ixazomib for 96 hours in media with or without 1 mM pyruvate. Growth rates are relative to the untreated condition, n=6 per condition. **(H)** Relative growth rate of A549 cells treated with or without 0.5 mM mesoxalate and cotreated with increasing doses of bortezomib in media containing 1 mM pyruvate for 96 hours. Growth rates are relative to the untreated condition, n=6 per condition. **(I)** Relative growth rate of A549 treated with or without 1 mM phenylpyruvate and cotreated with increasing doses of carfilzomib for 96 hours. Growth rates are relative to the untreated condition, n=6 per condition. **(J)** Relative growth rate of A549 treated with or without 0.5 mM mesoxalate and cotreated with varying doses of carfilzomib for 96 hours. Growth rates are relative to the untreated condition, n=6 per condition. For all panels values represent means, error bars are SEM. Relative growth rate curves were fitted with a nonlinear variable slope (four parameters) regression model. Statistical significance was assessed by an unpaired two-tailed student’s t-test (C). **** p<0.0001.

**Extended Data Figure 6.**
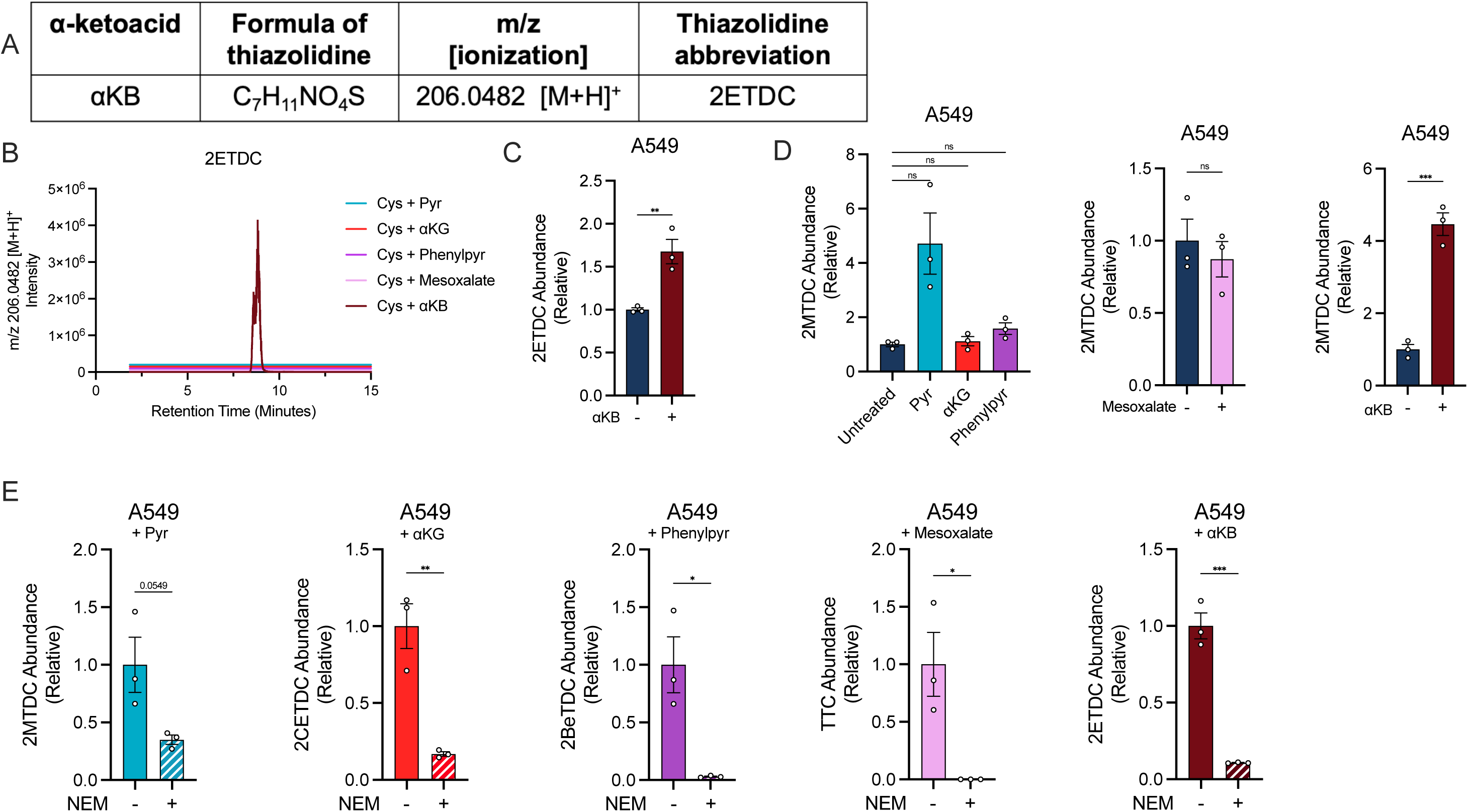
α-ketoacids react with cysteine to form specific thiazolidines in a reversible manner. **(A)** Table of predicted thiazolidine between cysteine and the α-ketoacid αKB, its chemical formula, its predicted m/z and ionization and abbreviation: CYS-αKB <u>2</u>-<u>e</u>thyl- 1,3-thiazolidine-2,4-<u>dic</u>arboxylate (2ETDC). **(B)** LC-MS ion chromatogram filtered for the predicted thiazolidine between cysteine and αKB. Purified chemical standards of CYS were incubated with or without each of the α-ketoacids overnight at 4**°**C. **(C)** Relative abundance of 2ETDC as measured by LC-MS in A549 cells treated with or without αKB for 1.5 hours. Abundances are relative ion counts to untreated cells, n=3 per condition. **(D)** Relative abundance of 2MTDC as measured by LC-MS in A549 cells treated with or without various α-ketoacids for 1.5 hours, n=3 replicate wells per condition. **(E)** Relative abundance of 2MTDC, 2CETDC, 2BeTDC, TTC, or 2ETDC in A549 cells extracted with or without the conjugating agent N-ethylmaleimide (NEM), n=3 per condition. Abundances are relative ion counts to the methanol-extracted samples. For all panels values represent means, error bars are SEM. Statistical significance was assessed with a two-tailed unpaired student’s t-test (C, D (middle, right), E) or a unpaired two-tailed student’s t-test with Bonferroni correction (D, left). *p < 0.05, **p < 0.01, ***p < 0.001, ns = not significant.

**Extended Data Figure 7.**
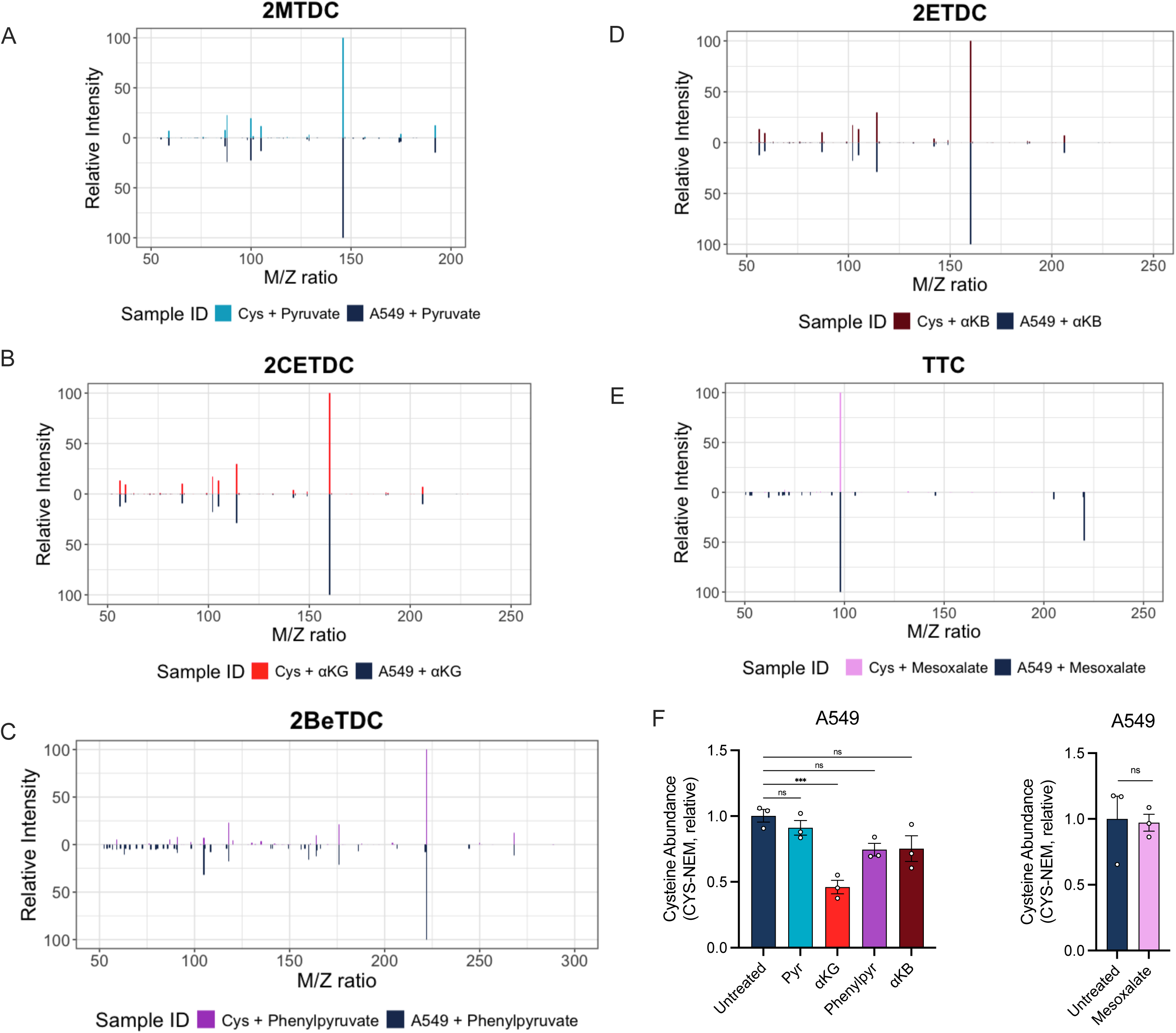
MS/MS fragmentation pattern for thiazolidine fates resulting from the reactions between α-ketoacids and cysteine. **(A)** Tandem mass spectrometry (MS/MS) fragmentation pattern for the thiazolidine fate, 2MTDC, generated in a cell-free system by combining pyruvate with CYS compared to the corresponding analyte extracted from A549 cells. **(B)** Tandem mass spectrometry (MS/MS) fragmentation pattern for the thiazolidine fate, 2CETDC, generated in a cell-free system by combining αKG with CYS compared to the corresponding analyte extracted from A549 cells. **(C)** Tandem mass spectrometry MS/MS fragmentation pattern for the thiazolidine fate, 2BeTDC, generated in a cell-free system by combining phenylpyruvate with CYS compared to the corresponding analyte extracted from A549 cells. **(D)** Tandem mass spectrometry (MS/MS) fragmentation pattern for the thiazolidine fate, 2ETDC, generated in a cell-free system by combining αKB with CYS compared to the corresponding analyte extracted from A549 cells. **(E)** Tandem mass spectrometry MS/MS fragmentation pattern for the thiazolidine fate, TTC, generated in a cell-free system by combining mesoxalate with CYS compared to the corresponding analyte extracted from A549 cells. **(F)** Relative abundance of total intracellular CYS, measured as CYS-NEM, from A549 cells with various α-ketoacids for 1.5 hours, then extracted with NEM and measured by LC-MS metabolomics. Abundances are relative ion counts to untreated cells. n=3 replicate wells per condition. For all panels values represent means, error bars are SEM. Statistical significance was assessed with a one-way ANOVA with Sidak’s correction for multiple comparisons (F, left) or an unpaired two-tailed student’s t-test (F, right). ***p < 0.001, ns = not significant.

**Extended Data Figure 8.**
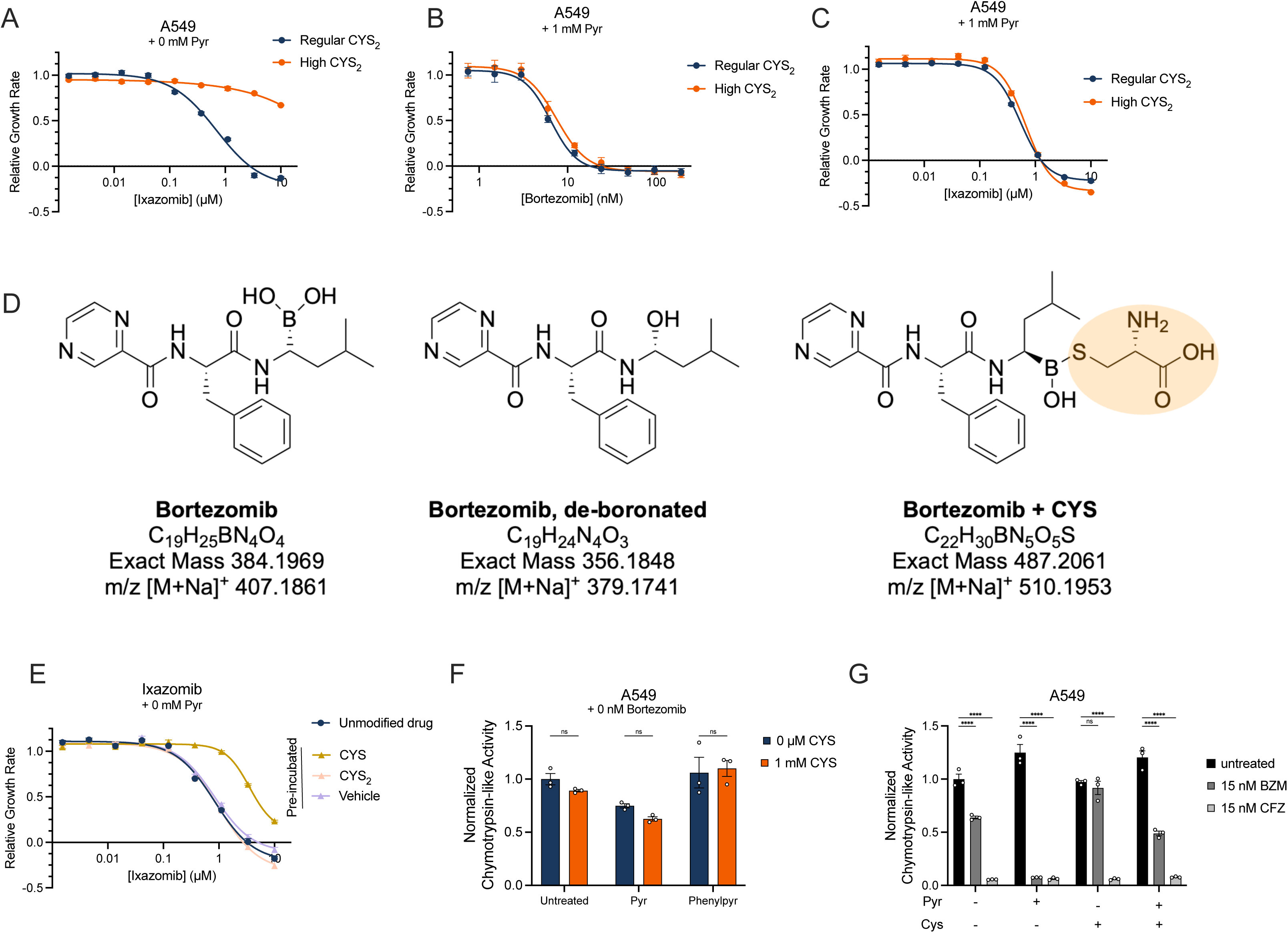
Cysteine directly inactivates boronic acid proteasome inhibitors to reduce their cytotoxicity. **(A)** Relative growth rate of A549 cells upon treatment with 200 μM CYS_2_ or 700 μM CYS_2_ and cotreatment with varying doses of ixazomib as measured by SRB. Growth rates are relative to the 0 nM ixazomib treated wells for each treatment condition. n=6 replicate wells per condition. **(B)** Relative growth rate of A549 cells upon treatment with 200 μM CYS_2_ or 700 μM CYS_2_ and 1 mM pyruvate and cotreatment with varying doses of bortezomib as measured by SRB. Growth rates are relative to the 0 nM bortezomib treated wells for each treatment condition. n=6 replicate wells per condition. **(C)** Relative growth rate of A549 cells upon treatment with 200 μM CYS_2_ or 700 μM CYS_2_ and 1 mM pyruvate and cotreatment with varying doses of ixazomib as measured by SRB. Growth rates are relative to the 0 nM ixazomib treated wells for each treatment condition. n=6 replicate wells per condition. **(D)** Chemical structures of proposed bortezomib fates including name of fate, exact mass, and Na LC-MS adduct. **(E)** Relative growth rate of A549 cells upon treatment with unmodified ixazomib or ixazomib that had been previously incubated with CYS, CYS_2_, or PBS overnight at 4**°**C as measured by SRB. n=6 replicate wells per condition. **(F)** Chymotrypsin-like proteasome activity was measured in A549 cells treated with 1 mM pyruvate or 1 mM phenylpyruvate with and without supplementation of 1 mM cysteine. n=3 replicate wells per condition. **(G)** Chymotrypsin-like proteasome activity was measured in A549 cells treated with and without 1 mM pyruvate, 1 mM cysteine, 15 nM bortezomib, or 15 nM carfilzomib, as indicated. n=3 replicate wells per condition. For all panels values represent means, error bars are SEM. Relative growth rate curves were fitted with a nonlinear variable slope (four parameters) regression model. Statistical significance was assessed by a two-way ANOVA with Sidak’s test for multiple comparisons (F, G). ns = not significant, ****p < 0.0001.

